# Cholinergic neuronal responses to probabilistic outcome-predicting stimuli follow a weighed, unsigned prediction error model and anticipate behavioral responses

**DOI:** 10.1101/2022.07.05.498795

**Authors:** Panna Hegedüs, Katalin Sviatkó, Bálint Király, Sergio Martínez-Bellver, Balázs Hangya

## Abstract

Basal forebrain cholinergic neurons (BFCNs) play an important role in associative learning, suggesting that BFCNs may participate in processing sensory stimuli that predict future outcomes. However, little is known about how BFCNs respond to outcome-predictive sensory cues and the impact of outcome probabilities on BFCN responses has not been explored. Therefore, we performed bulk calcium imaging and recorded spiking output of identified cholinergic neurons from the basal forebrain of mice performing a probabilistic Pavlovian cued outcome task that allowed us to control the predictive strength of cue stimuli. BFCNs responded strongly to sensory cues predicting likely reward, while little response was observed for cues that were rarely paired with reward. Reward delivery led to the activation of BFCNs, with less expected rewards eliciting a stronger response, while air puff punishments also evoked positive-going responses from BFCNs. We propose that BFCNs differentially weigh predictions of positive and negative reinforcement, reflecting divergent relative salience of forecasting appetitive and aversive outcomes, in accordance with a simple reinforcement learning model of a weighed, unsigned prediction error. Finally, the extent of cholinergic activation after cue stimuli predicted subsequent decision speed, suggesting that the expectation-gated cholinergic firing is instructive to reward-seeking behaviors.

## Introduction

Cholinergic neurons of the basal forebrain are important for associative learning. This idea is supported by the selective cholinergic cell loss that parallels cognitive decline in Alzheimer’s disease patients (Whitehouse et al., 1982; Arendt and Bigl, 1986). While lesion and pharmacology studies were confirmative (Everitt and Robbins, 1997; Wrenn and Wiley, 1998; Hasselmo and Sarter, 2011), they cannot solve how BFCNs exert their control over learning. To address the mechanisms of the contribution of BFCNs to associative learning, it is important to investigate the behavioral correlates of BFCN activity at temporal resolutions comparable to the time scales of behaviorally relevant events animals and humans encounter (Sviatkó and Hangya, 2017; Solari et al., 2018). This has only become possible recently, enabled by the development of new optogenetic and imaging tools (Lovett-Barron et al., 2014; Hangya et al., 2015; Harrison et al., 2016; Guo et al., 2019).

Selective cholinergic lesions of the basal forebrain were shown to impair learning in rodents (Berger-Sweeney et al., 1994; McGaughy et al., 2002, 2005; Bailey et al., 2003; Conner et al., 2003; Dalley et al., 2004; Ross et al., 2005) and monkeys (Fine et al., 1997), and lesions of the basal forebrain in consequence of aneurysm rupture of the anterior cerebral or anterior communicating artery leads to severe learning impairments in humans (Damasio et al., 1985). Previous studies of the basal forebrain have proposed that responses to behaviorally salient stimuli of cholinergic and/or non-cholinergic basal forebrain neurons may underlie the involvement of the basal forebrain in learning (Lin and Nicolelis, 2008; Hangya et al., 2015; Harrison et al., 2016; Guo et al., 2019; Crouse et al., 2020). Specifically, cholinergic activation may lead to increased cortical acetylcholine release that induces plastic changes in sensory responses (Kilgard and Merzenich, 1998; Froemke et al., 2007). A recent study connected the above pieces of evidences by bulk imaging of BFCNs during auditory fear learning (Guo et al., 2019). However, it is not yet known how BFCNs process sensory cues with different predictive features during learning, which could serve as a basis for differential behavioral responses to sensory events that forecast distinct outcomes. Therefore, a comprehensive model of cholinergic neuronal responses that subserve associative learning is also lacking. We set out to fill this knowledge gap by recording cholinergic activity in a probabilistic Pavlovian cued outcome task, which allowed us to directly control outcome probabilities and cue-outcome contingencies during learning (Hegedüs et al., 2021b). Of note, reward expectation can also be manipulated by reward size (Eshel et al., 2016; Stephenson-Jones et al., 2020). However, since we hypothesized that BFCNs are sensitive to the outcome probabilities, we chose to manipulate reward probability instead, despite that this is harder to learn, as animals have to integrate over multiple trials to infer differences in probability, whereas reward size can be learned from a single trial (Hegedüs et al., 2021a).

We imaged the bulk calcium responses of BFCNs using fiber photometry (Kim et al., 2016) and recorded the activity of identified basal forebrain cholinergic neurons while mice were performing a head-fixed auditory probabilistic Pavlovian cued outcome task (Hegedüs et al., 2021b). BFCNs were activated by outcome-predicting stimuli, as well as delivery of reinforcement. Reward-predicting stimuli activated cholinergic neurons differentially in correlation with the likelihood of future reward, and subsequent reaction times were predicted by the level of this activation. BFCNs also showed stronger activation after unexpected compared to expected rewards. We show that these findings can be explained by a behavioral model of a weighed, unsigned prediction error, in which outcomes of opposite valence are differentially scaled by the animals. We did not observe robust firing rate changes of BFCNs following omissions of reinforcement, suggesting that the BFCN responses we observed were largely driven by sensory stimuli. Thus, these results suggest that the central cholinergic system broadcasts a stimulus-driven, valence-weighed, unsigned prediction error signal that can instruct associative learning.

## Results

We trained mice (n = 11) on a head-fixed probabilistic Pavlovian cued outcome task (Fig. 1A) (Hegedüs et al., 2021a, 2021b). During this associative learning task, two tones of different pitch (conditioned stimuli) predicted either water reward with 80% chance (10% punishment, 10% omission, ‘likely reward’ cue) or a puff of air on the face with 65% probability (25% reward, 10% omission, ‘unlikely reward’ cue; the contingencies reflect careful calibration to keep mice motivated for the task). Based on the cue that preceded behavioral feedback (unconditioned stimuli), both rewards and punishments could be either expected, or surprising. Mice learned this task, indicated by performing significantly more anticipatory licks after ‘likely reward’ cues (Fig. 1B-E).

**Fig 1.**
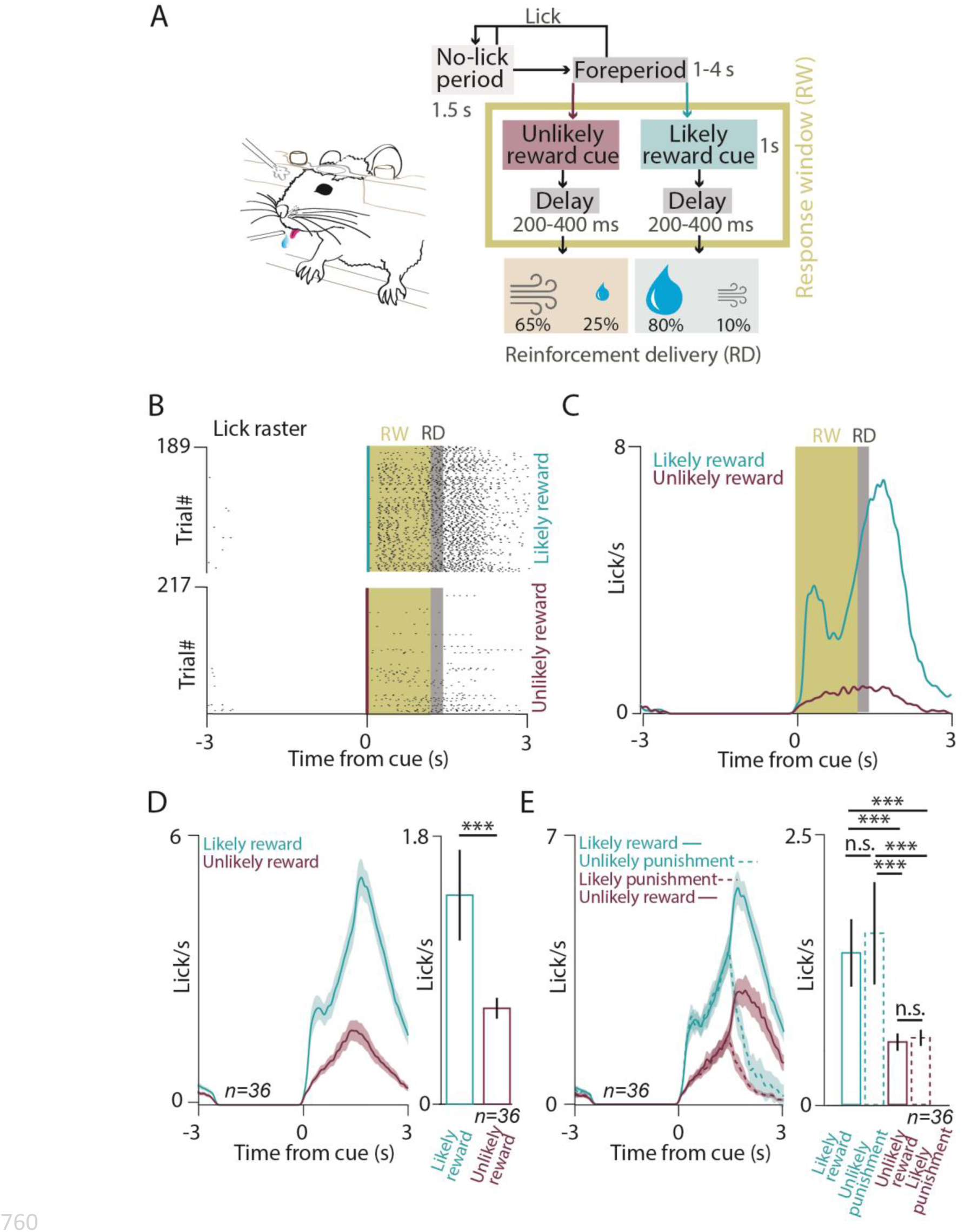
Animals were trained on a probabilistic Pavlovian conditioning task. (A) Schematic illustration of the behavioral training and block diagram of the task. A variable foreperiod, in which the mouse was not allowed to lick, was followed by the presentation of one of two pure tones of well-separated pitch, which predicted reward, punishment or nothing with different contingencies (‘likely reward’ and ‘unlikely reward’ cues). (B) Raster plot of lick responses to the cues predicting likely reward (top) and unlikely reward (bottom) from an example session. Yellow shading, response window (RW); gray shading, reinforcement delivery (RD). (C) Peri-event time histograms (PETHs) of lick responses aligned to cue onset in the same session. (D) Left, average PETHs of lick responses of all sessions of all animals (n = 36 sessions); Right; statistical comparison of anticipatory lick rates in the RW in likely reward and unlikely reward trials (median ± SE of median, n = 36 sessions, p = 8.7697 x 10^-7^, Wilcoxon signed-rank test; ***, p < 0.001; n.s., p > 0.05). (E) Left, average PETHs of lick responses of all sessions of all animals (n = 36 sessions), partitioned based on four possible outcomes: expected or surprising reward, expected or surprising punishment. Right, statistical comparison of anticipatory lick rates in the RW in the four possible outcomes (median ± SE of median, n = 36 sessions, from top to bottom, p = 1.4131 x 10^-6^, p = 2.6341 x 10^-6^, p = 0.8628, p = 6.8863 x 10^-7^, p = 1.2065 x 10^-6^, p = 0.9687, Wilcoxon signed-rank test; ***, p < 0.001; n.s., p > 0.05).

We expressed GCaMP6s in BFCNs of the horizontal nucleus of the diagonal band of Broca (HDB) by injecting AAV2/9.CAG.Flex.GCAMP6s.WPRE.SV40 in ChAT-Cre mice (n = 7), expressing Cre-recombinase driven by the choline-acetyltransferase promoter selectively in cholinergic neurons (Higley et al., 2011; Eggermann et al., 2014; Hangya et al., 2015), and implanted them with an optic fiber in the HDB. We performed bulk calcium imaging of HDB BFCNs while mice were performing the probabilistic Pavlovian task (Fig. 2A-C, Fig. S1). An excitation isosbestic wavelength of GCaMP was used to correct for non-calcium-dependent changes in fluorescence (e.g. bleaching and potential movement artifacts) (Lerner et al., 2015). We first asked whether BFCNs as a population responded to auditory cue stimuli that predicted outcomes with different contingencies. Fluorescent dff responses were aligned to cue presentations, revealing BFCN population calcium responses to outcome-predicting cue stimuli (Fig2. D-E). These responses were significantly larger for cues that predicted ‘likely reward’ (and ‘unlikely punishment’) compared to cues predicting ‘likely punishment’ (and ‘unlikely reward’; Fig2. D-E, p = 0.00029, Wilcoxon signed-rank test, n = 17 sessions). Based on published results of us and others (Lovett-Barron et al., 2014; Hangya et al., 2015; Harrison et al., 2016; Guo et al., 2019), we expected BFCN calcium responses following the delivery of rewards and punishments as well. Indeed, when dff recordings were aligned to reinforcement, we observed robust cholinergic population responses to both water reward and air puff punishment (Fig. 2F-G). Moreover, we found that BFCN responses to surprising rewards significantly exceeded those to expected rewards, although the observed difference was less than that of the cue responses (Fig. 2F, p = 0.0129, Wilcoxon signed-rank test, n = 17 sessions). We did not find a significant difference between BFCN calcium responses to surprising vs. expected punishments (Fig. 2G, p = 0.0684, Wilcoxon signed-rank test, n = 17 sessions).

**Fig 2.**
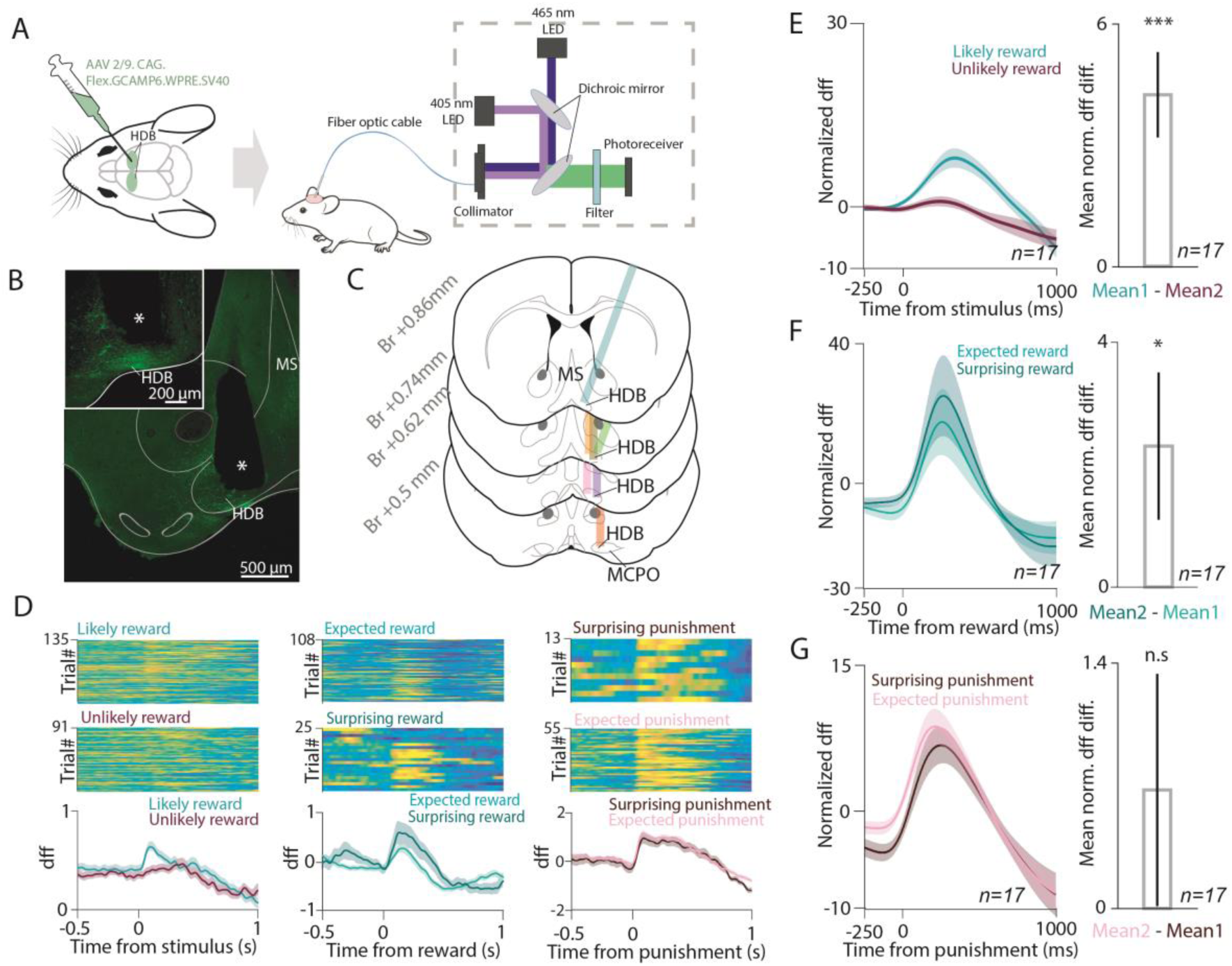
BFCN population responses to conditioned and unconditioned stimuli. (A) Schematic diagram of bulk calcium-imaging of HDB BFCNs in behaving mice. (B) Histological reconstruction of the optic fiber track in the HDB (white asterisk). (C) Optical fiber locations in all imaged mice (n = 7). Br, antero-posterior distance from Bregma. (D) Example session of bulk calcium imaging of cholinergic neurons from the HDB. Left, dff signals were aligned to outcome-predicting conditioned stimuli. Top, trials with likely reward (unlikely punishment) cue; middle, trials with unlikely reward (likely punishment) cue; bottom, PETH. Middle, dff signals were aligned to reward delivery. Top, trials with expected reward; middle, trials with surprising reward; bottom, PETH. Right, dff signals were aligned to air puff punishments. Top, trials with surprising punishment. Middle, trials with expected punishment. Bottom, PETH. (E) Left, average PETH of Z-scored dff aligned to outcome-predicting conditioned stimuli (n = 17 sessions). Right, bar graph of average normalized difference in dff after likely reward and unlikely reward cues. Median ± SE of median, ***, p < 0.001, p = 0.00029, Wilcoxon signed-rank test. (F) Left, average PETH of Z-scored dff aligned to expected and surprising reward (n = 17 sessions). Right, bar graph of average normalized difference in dff after surprising and expected reward. Median ± SE of median, *, p < 0.05, p = 0.0129, Wilcoxon signed-rank test. (G) Left, average PETH of Z-scored dff aligned to expected and surprising punishment (n = 17 sessions). Right, bar graph of average normalized difference in dff after expected and surprising punishment. Median ± SE of median, n.s., p > 0.05, p = 0.0684, Wilcoxon signed-rank test.

Does the spiking of individual BFCNs show similar differential responses to conditioned and unconditioned stimuli according to different outcome expectations? We estimated that a sample of 14-20 identified BFCNs is sufficient to answer such a question with 80% statistical power (assuming a 30-40% firing rate change corresponding to 0.3-0.4 predicted effect size, detectable in 60% of recorded neurons; full procedure available at https://github.com/hangyabalazs/statistical-power; Fig. 3A). We expressed channelrhodopsin in BFCNs by injecting AAV.2.5.EF1a.DiO.hChR2(H134R).eYFP.WPRE.hGh in the basal forebrain of ChAT-Cre mice (n = 4) and implanted them with eight moveable tetrode electrodes and an optic fiber (Fig. 3B-C), in order to optogenetically tag BFCNs in mice performing the probabilistic Pavlovian task (Hangya et al., 2015; Guo et al., 2019). We recorded 25 optogenetically identified, ChAT-expressing BFCNs (p < 0.01, stimulus-associated spike latency test (Kvitsiani et al., 2013)) in task-performing mice (Fig. 3D-G; Fig. S2-3). Careful post-hoc histological reconstruction of the location of the recording electrodes showed that 21/25 neurons were recorded from HDB, while the remaining 4/25 neurons were in the medial septum (n = 2) and the ventral pallidum (n = 2; Fig. 3D). Since these neurons exhibited similar responses to conditioned and unconditioned stimuli, they were treated as a single data set for this study; nevertheless, restricting data analyses to the HDB cholinergic neurons yielded similar results.

**Figure 3.**
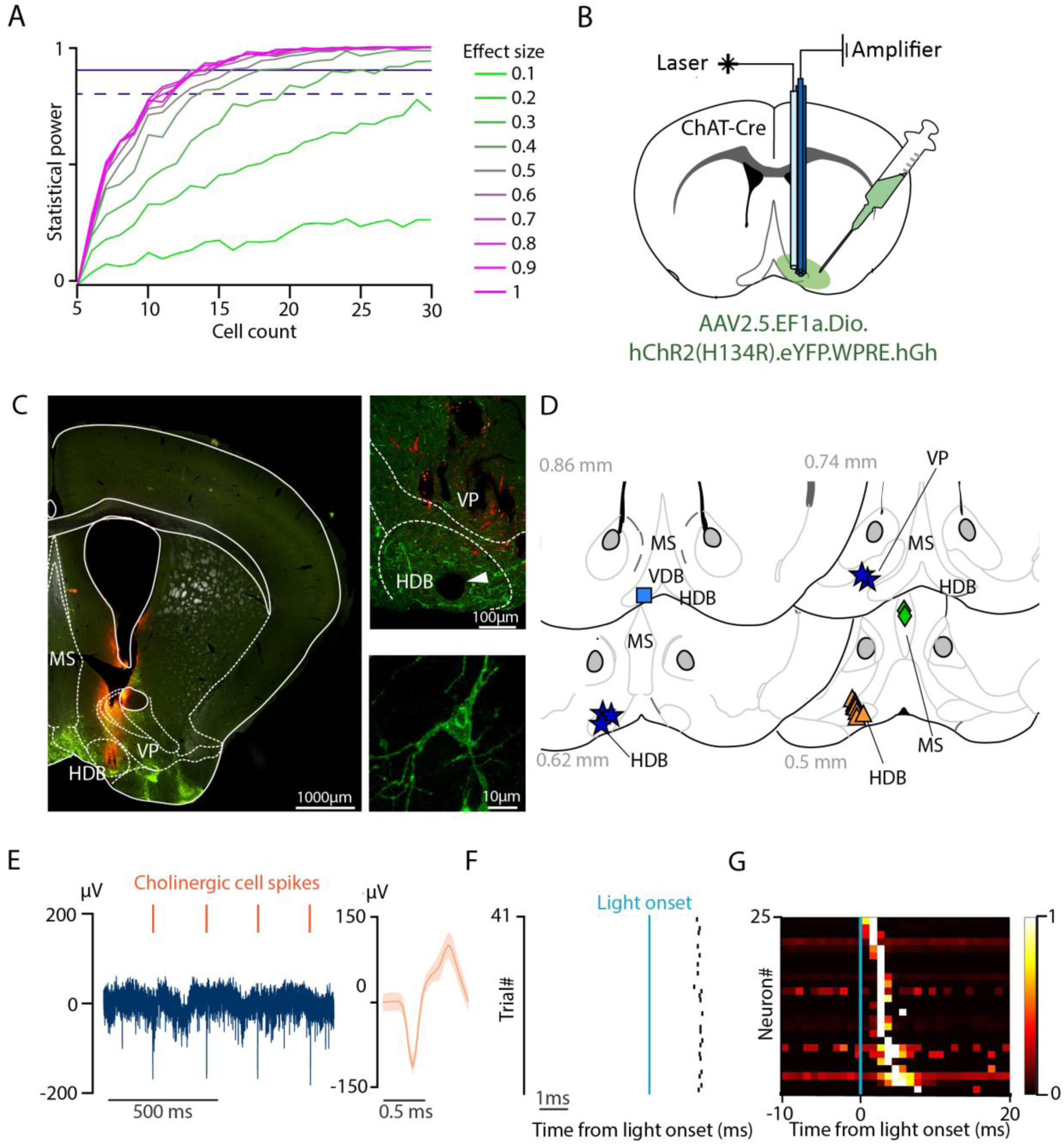
Optogenetic identification of basal forebrain cholinergic neurons during probabilistic Pavlovian conditioning. (A) Statistical power as a function of cell count at different expected effect sizes. (B) Schematic drawing of optogenetic tagging. ChAT-Cre mice were injected with AAV2/5. EF1a.Dio.hChR2(H134R)-eYFP.WPRE.hGH. Eight moveable tetrodes were implanted in the HDB along with an optic fiber. (C) Left, coronal section from a ChAT-Cre mouse showing the distribution of cholinergic neurons (eYFP, green) and the tetrode track (DiI, red). Top right, magnified view of the HDB. The white arrowhead points to the electrolytic lesion marking the tetrode tips. Bottom right, confocal image of a cholinergic neuron from the target area. (D) Reconstructed localization of all identified cholinergic neurons. Different markers correspond to individual mice. (E) Left, raw extracellular recording of an identified cholinergic neuron. Right, average waveform of the example cholinergic neuron. Orange marks above the recording indicate the cholinergic spikes. (F) Raster plot of an example cholinergic neuron showing short latency responses to 1 ms blue laser pulses. (G) Color-coded PETH of all identified cholinergic neurons aligned to laser pulse onset, sorted by response latency (black, no spikes; white, high firing rate).

We first asked whether individual BFCNs show spiking responses to auditory cue stimuli that predict outcomes with different probabilities. To address this, we aligned BFCN spikes to cue onset and examined raster plots and peri-event time histograms (PETH) of individual BFCNs (see Fig. S4 for a schematic representation of the analysis). We found that BFCNs responded to both auditory cues, with a median peak latency of 133.5 ms for the ‘likely reward’ cue and 422 ms for the ‘unlikely reward’ cue (Fig. 4A-B, Fig. S5A; interquartile range, 44.5-231 ms and 273-573.5 ms for the two cue types). To cover both peaks, we chose a 500 ms response window (C500), in which we compared BFCN responses to conditioned cue stimuli based on whether they signaled high or low probability of future reward. BFCNs showed 151% stronger responses to the ‘likely reward’ cues based on a comparison of PETH peak responses in the C500 window (p = 0.0008, Wilcoxon signed-rank test; Fig. 4C; including n = 14 neurons where mice encountered >10 surprising reward trials; see Fig. S6 for all n = 25 neurons), which we also confirmed by spike-number-based statistics (p = 0.00061, Wilcoxon signed rank test on BFCN firing rates in the C500 window; Fig. 4C). Thus, BFCNs responded more to sensory stimuli that signaled high probability of reward.

**Figure 4.**
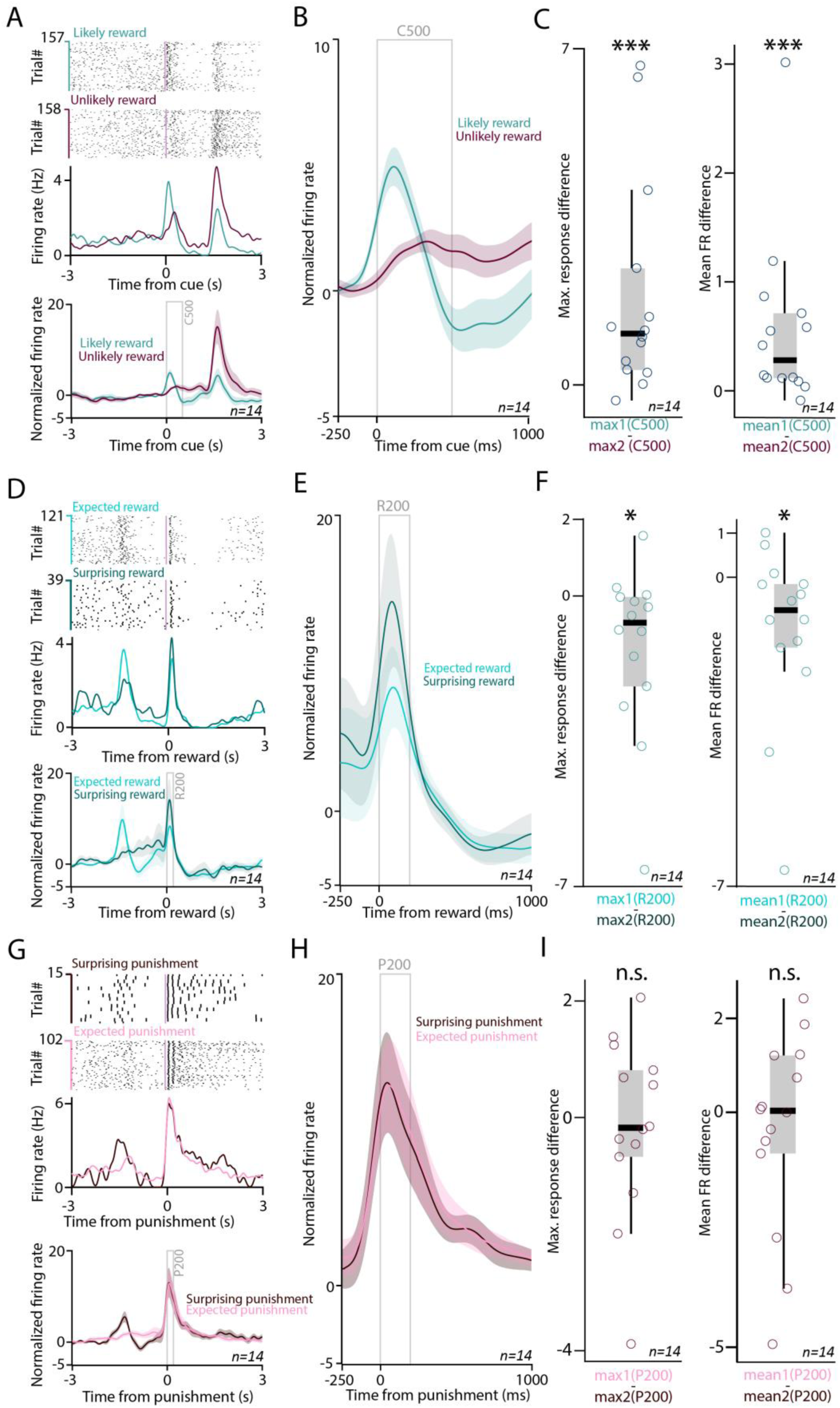
Cholinergic neurons respond more to reward-predicting cues and surprising reward. (A) Top, Raster plots (top) and PETHs (bottom) of an example BFCNs aligned to cue onset, separately for the cues predicting likely reward / unlikely punishment (turquoise) vs. unlikely reward / likely punishment (purple). Bottom, average cue-aligned PETHs of identified BFCNs with >10 surprising reward trials (errorshade, mean ± SE; n = 14; see Fig. S6 for all n = 25 neurons). (B) Average cue-aligned PETHs of identified BFCNs enlarged around cue presentations. (C) Left, difference in peak response after cues predicting likely reward and those predicting unlikely reward. ***, p < 0.001, p = 0.0008, Wilcoxon signed-rank test, n = 14. Right, Difference in mean firing rate after cues predicting likely reward and those predicting unlikely reward. ***, p < 0.001, p = 0.00061, Wilcoxon signed-rank test, n = 14. (D) Top, raster plots (top) and PETHs (bottom) of the same example BFCN as in A, aligned to reward delivery, separately for rewards after the cue predicting likely reward (light turquoise, expected reward) vs. after the cue predicting unlikely reward (dark turquoise, surprising reward). Bottom, average reward-aligned PETHs of identified BFCNs with >10 surprising reward trials (errorshade, mean ± SE; n = 14; see Fig. S6 for all n = 25 neurons). (E) Average reward-aligned PETHs of identified BFCNs enlarged around reward delivery times. (F) Left, difference in peak response after expected and surprising rewards. *, p < 0.05, p = 0.0245, Wilcoxon signed-rank test, n = 14. Right, difference in mean firing rate after expected and surprising rewards. *, p < 0.05, p = 0.02026, Wilcoxon signed-rank test, n = 14. (G) Top, raster plots (top) and PETHs (bottom) of the same example BFCNs as in A and D, aligned to punishment delivery, separately for punishments after the cue predicting likely reward (dark purple, surprising punishment) vs. after the cue predicting unlikely reward (light purple, expected punishment). Right, average punishment-aligned PETHs of identified BFCNs enlarged around punishment delivery times. (H) Average punishment-aligned PETHs of identified BFCNs with >10 surprising reward trials (errorshade, mean ± SE; n = 14; see Fig. S6 for all n = 25 neurons). (I) Left, difference in peak response after expected and surprising punishments. n.s., p > 0.05, p = 0.7869, Wilcoxon signed-rank test, n = 14. Right, difference in mean firing rate after expected and surprising punishments. n.s., p > 0.05, p = 0.8393, Wilcoxon signed-rank test, n = 14.

Next, we tested whether individual BFCNs responded to the delivery of reward during Pavlovian conditioning, and whether this response depended on previous expectations about reward likelihoods conveyed by the two auditory cues. Therefore, we aligned the spike times of the same BFCNs to the time of reward delivery, again examining raster plots and PETHs (Fig. 4D-E). We found that reward also elicited large BFCN responses, with a median peak latency of 86.5 ms and 82.7 ms for expected and surprising reward, respectively (Fig. S5B; interquartile range, 78.13-100.25 ms and 54.5-92.5 ms for expected and surprising reward). To compare BFCN responses to expected vs. surprising rewards, we defined a 200 ms response window after reward delivery based on the above latency measurements (R200). We found that rewards that were less expected lead to significantly stronger cholinergic firing (69.3%, p = 0.0245, Wilcoxon signed-rank test on R200 response peaks; Fig. 4F), also confirmed by firing rate comparison (p = 0.02026, Wilcoxon signed-rank test on BFCN firing rates in the R200 window; Fig. 4F). These findings showed that BFCN responses were modulated by the expectation of reward.

We took a similar approach to investigate BFCN responses to the delivery of air puff punishments. BFCNs also responded with firing rate increase to punishment, with remarkably short peak latencies (Fig. 4G-H, Fig. S6C; median and interquartile range, 24.5 ms and 15.5 - 36 ms for surprising punishment and 24 ms and 15.5 - 32 ms for expected punishment), confirming previous results (Lovett-Barron et al., 2014; Hangya et al., 2015; Laszlovszky et al., 2020). When responses to surprising and expected punishments were directly compared in a 200 ms response window (P200), we did not find significant modulation by expectation (p = 0.7869, Wilcoxon signed-rank test on peak responses; Fig. 4I; p = 0.8393, Wilcoxon signed-rank test on firing rates). We did not detect significant firing rate changes in either direction after omissions (Fig. S7).

The above-demonstrated differential BFCN responses to conditioned and unconditioned stimuli that reflected outcome expectations was suggestive of prediction error coding (Schultz et al., 1997). Based on the positive-going BFCN responses following both reward and punishment, we assumed that BFCNs might represent an unsigned prediction error. If an unsigned prediction error scaled positive and negative values equally, then it would track the expectation of reinforcement irrespective of valence. Thus, it would predict identical responses to conditioned cue stimuli that foreshadow reinforcement with a fixed probability, only sensitive to the rate of reinforcement omissions. However, cholinergic neurons showed stronger responses after cues predicting likely reward compared with those predicting unlikely reward but likely punishment. Therefore, our results suggest that BFCNs assign different weights to expected positive and negative outcomes, potentially related to the difference in absolute subjective values of the reinforcers. We did not observe BFCN responses to reinforcement omissions, suggesting that BFCN responses are driven by sensory stimuli and thus a stimulus-driven, valence-weighed, unsigned prediction error model could explain BFCN spiking dynamics.

To test this, we implemented and fitted a simple three-parameter reinforcement learning (RL) model (Schultz et al., 1997; Kim et al., 2020) on cholinergic responses:

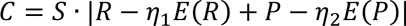

where *C* represented cholinergic response, *S* was a scaling parameter accounting for different mean firing rates of BFCNs, *R* and *P* were actual, while *E(R)* and *E(P)* were expected reward and punishment determined by task contingencies. To take the assumed difference in the relative sensitivity to water reward and air-puff punishment into account, we introduced two weight parameters, *η_1_* and *η_2_* (0 ≤ *η_1_*, *η_2_*≤ 1), which could control how much BFCN responses were influenced by the expectation of positive and negative outcomes, respectively. Taking the absolute value of the sum of reward and punishment prediction error terms ensured positive-going cholinergic responses irrespective of valence, thus resulting in a simple model of unsigned reward prediction error. We found that this model fitted BFCN firing rate changes in response to the different cues and reinforcers defined by the C500, R200 and P200 response windows well (Fig. 5A-C), and significantly better than a control model in which the modelled expectations did not match the task contingencies (p = 0.0014 for all n = 25 BFCNs recorded; p = 0.0037 when only HDB cholinergic neurons were tested; Wilcoxon signed-rank test on the maximum likelihoods of the models; see Methods).

**Figure 5.**
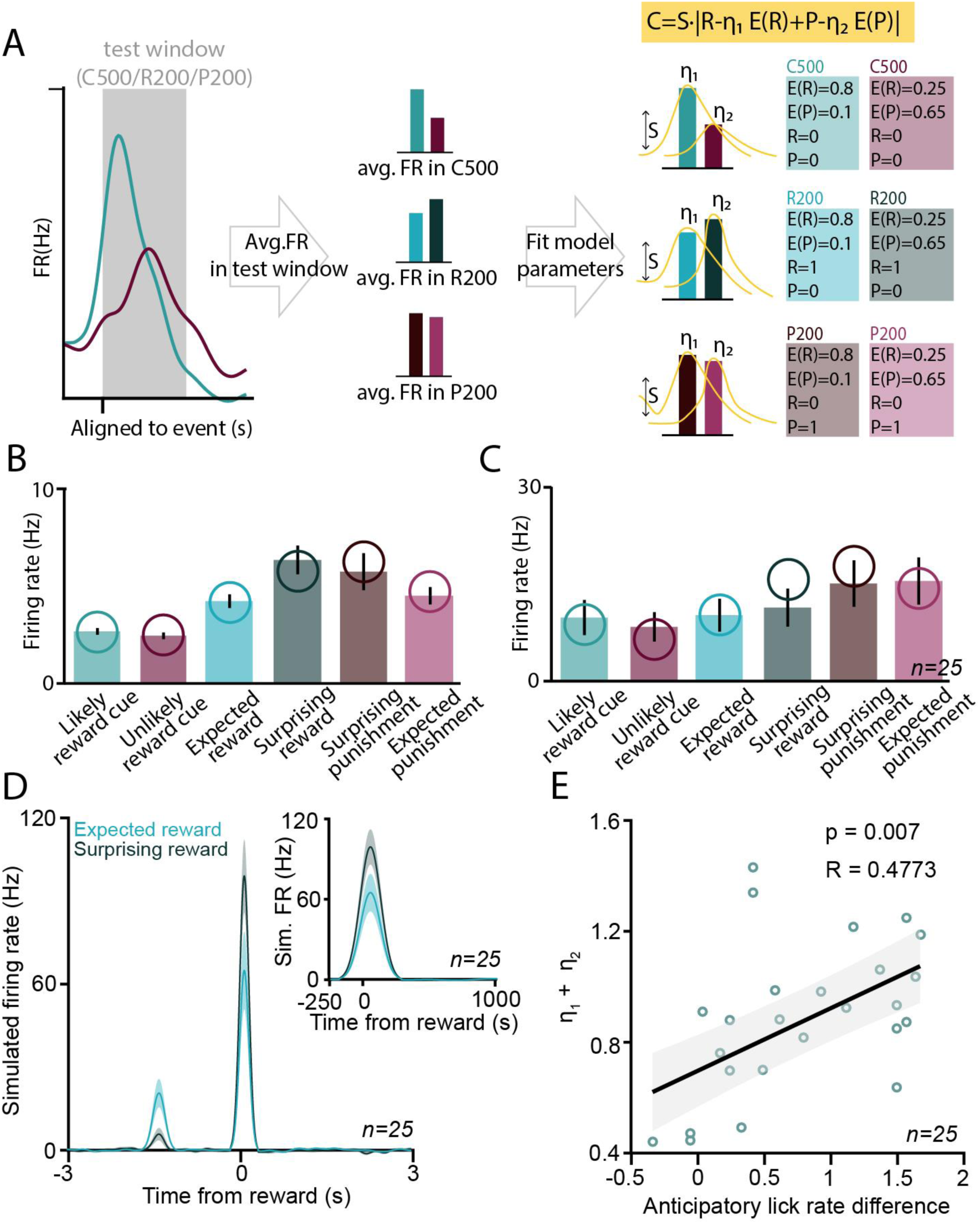
Cholinergic responses are explained by a reinforcement learning model of stimulus-driven, valence-weighed, unsigned prediction error. (A) Schematic illustration of reinforcement learning model fitting on cholinergic neuronal data. Average FR values were fit by a three-parameter RL model incorporating task contingencies. (B) Firing rates of an example BFCN in 500 ms response windows after cue presentations and 200 ms response windows after reward or punishment, separated by trial type. Bar graphs represent mean ± SE over trials. Hypothetical firing rates corresponding to a best-fit RL model are overlaid, indicated by open circles. (C) Average firing rates of all identified BFCNs (n = 25) in the same response windows. Bar graphs represent mean ± SE over neurons. Average modelled firing rates are indicated by open circles. (D) Spike responses were simulated based on the best-fit RL models for each BFCN (see Methods). PETHs were calculated the same way as for the real data and averaged over the modelled responses (n = 25). (E) The sum of the two model parameters that controls the differential sensitivity to reward and punishment expectations (*η_1_ + η_2_*) was correlated with the difference in anticipatory lick rate after likely reward vs. unlikely reward predicting cues (R = 0.4773, Pearson’s correlation coefficient; p = 0.007, linear regression, F-test).

We next simulated spike trains of individual BFCNs based on the best-fit RL models. Baseline firing was modelled by a Poisson process with a frequency matched to the baseline firing rate of the modelled BFCN, and simulated firing responses were added according to Gaussian distributions with a fixed delay after cue and reinforcement events, where the number of added spikes was determined by the best-fit RL model for each BFCN. When applying the same analyses on simulated spike trains as for the real data, we found that simulated PETHs qualitatively reproduced BFCN responses to cues and rewards (Fig. 5D). These results further strengthen that the BFCN responses we observed are consistent with the representation of a weighed, unsigned prediction error.

The best-fit *η_1_* values were significantly larger than the best-fit *η_2_* values, demonstrating stronger sensitivity of BFCN responses to reward than to punishment expectations (p = 0.0001, Wilcoxon signed-rank test). These parameters might reflect potential differences in the internal valuation of water reward and air-puff punishment across animals and recording days, but also different sensitivity to reward expectation of individual BFCNs. We hypothesized that they reflect behavioral variability rather than heterogeneity across neurons, which would imply that these parameters show consistency within recording sessions and within individual mice. Indeed, we found smaller within-than across-mice differences in best fit *η_1_* parameters (p = 0.002, Mann-Whitney U test), and smaller within-than across-session differences in best fit *η_2_* parameters (p = 0.047, Mann-Whitney U test; n = 25; Fig. S8). This suggests that best fit scaling parameters for outcome expectations reflect inter-individual and/or behavioral differences, rather than differential sensitivity of individual BFCNs.

The perceived reward and punishment prediction errors are controlled by *η_1_* and *η_2_* in our model; therefore, they together determine the size of the unsigned outcome prediction error represented by BFCNs. If this can drive approach behaviors as previous studies suggested (Lin and Nicolelis, 2008; Avila and Lin, 2014; Ahrens et al., 2018; Zhang et al., 2019), then we would expect that the animals’ anticipatory licking behavior correlates with these model parameters. Indeed, we found that *η_1_* as well as the sum of the two parameters (*η_1_ + η_2_*), characterizing the cholinergic neurons’ sensitivity to momentary outcome prediction, correlates well with behavioral cue differentiation as indexed by anticipatory lick rate difference (p = 0.012, R = 0.52 and p = 0.056, R = 0.33 for *η_1_* and *η_2_*, respectively; p = 0.007, R = 0.48 for *η_1_+ η_2_*; n = 25; Fig. 5E and Fig. S8; p = 0.0013 and R = 0.78 when calculated for the n = 14 neurons with >10 surprising reward trials; Pearson’s correlation coefficient, linear regression and one-sided F-test).

The correlation of model parameters quantifying the animals’ sensitivity to outcome expectations with behavioral performance prompted us to further assess whether BFCN responses could predict animal behavior. BFCN responses to outcome predicting cues consistently preceded the animals’ first licks (Fig. 6A-B). When we aligned cholinergic spikes to the last lick before the foreperiod during which mice were not allowed to lick, cholinergic activity peaked before licking with a similar time course as during cue-related licking activity (Fig. S9). These findings excluded that a potential ‘lick-driven’ cholinergic activity could confound the results and instead indicated that cholinergic activity had the potential to influence behavioral responses of mice performing the task. Indeed, we found that larger cholinergic cue responses were followed by faster reactions (p = 0.00073 and p = 0.05108 for ‘likely reward’ and ‘unlikely reward’ cues, respectively; Wilcoxon signed-rank test; Fig. 6C). In accordance, cholinergic cue responses were larger when mice were licking after the cue (p = 0.048 and p = 0.023 for ‘likely reward’ and ‘unlikely reward’ cues, respectively; Wilcoxon signed-rank test; Fig. 6D). Since lick responses can be taken as an indication of mice expecting reward, these results are consistent with cholinergic reward expectation coding. Next, we divided the trials into four quartiles according to mice’s reaction time after cue onset. In line with the above results, we found that faster lick responses were preceded by stronger cholinergic firing (Fig. 6E, p = 0.0314, one-way ANOVA). In sum, these results indicate that cue responses of BFCNs predict reaction times, suggesting that cholinergic outcome prediction coding affects behavioral responses.

**Figure 6.**
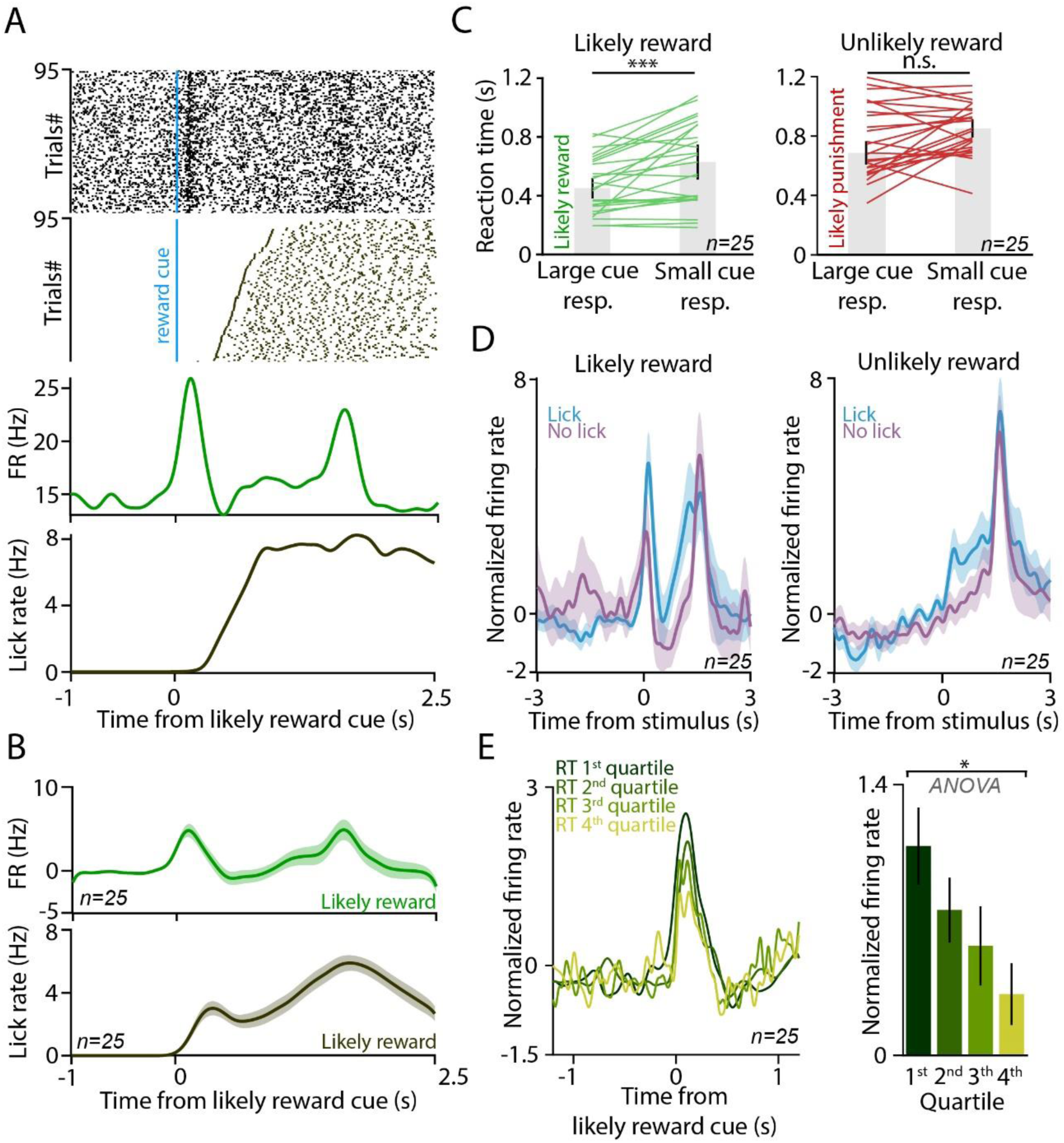
Cholinergic responses predict reaction time. (A) Top, spike raster of cholinergic firing (top) and lick responses of the animal (bottom) aligned to the likely reward cues from an example session. Bottom, corresponding PETH of the cholinergic response (top) and licking activity (bottom). (B) Average PETH of cholinergic firing (top) and lick response (bottom) after the reward-predicting cue (n = 25 BFCNs). (C) Reaction time after large and small cue responses to the likely reward- (left) and unlikely reward cue (right). Cue responses were divided by a median split. ***, p < 0.001, p = 0.00073 and n.s., p > 0.05, p = 0.05108, Wilcoxon signed-rank test, n = 25. (D) Average PETH of cholinergic responses to the likely reward and unlikely reward cue, separated based on the presence or absence of anticipatory lick response of the animal. (E) Stronger cholinergic response to the reward-predicting cue predicted faster reaction time. Left, average PETH of responses to the likely reward cue, partitioned to reaction time quartiles (1st quartile corresponds to shortest reaction time, in dark green). Right, bar graphs (mean ± SE) of the peak responses to the likely reward cue as a function of reaction time quartiles. *, p < 0.05, p = 0.0314, one-way ANOVA.

## Discussion

Cholinergic neurons of the basal forebrain respond to behaviorally salient events (Lovett-Barron et al., 2014; Hangya et al., 2015; Harrison et al., 2016; Teles-Grilo Ruivo et al., 2017; Guo et al., 2019; Crouse et al., 2020; Laszlovszky et al., 2020; Robert et al., 2021). To better understand the nature of these responses, we investigated whether activity patterns elicited by outcome-predictive stimuli and behavioral feedback are consistent with a prediction error hypothesis. By investigating the responses of BFCN populations using bulk calcium imaging and of individual BFCNs by optogenetic tagging in a probabilistic Pavlovian cued outcome task, we found that BFCNs showed strong activation after reward-predicting stimuli, and larger responses to surprising than to expected rewards. These results were consistent with a simple RL model of a stimulus-driven, valence-weighed, unsigned reward prediction error. The model also demonstrated that while BFCNs responded with firing rate increase to events of both positive and negative valence, they also reflected different behavioral sensitivity to positive and negative expectations. Finally, BFCN responses were found to likely influence behavioral performance, as mice showed faster responses after stronger cholinergic activation.

Temporal difference reinforcement learning (TDRL) models were successful in explaining the reward prediction errors represented by the dopaminergic system (Schultz et al., 1997; Gershman and Uchida, 2019; Kim et al., 2020). The presence of a reward response modulated by expectation and responsiveness to reward-predicting sensory stimuli suggested that cholinergic signals may also be related to prediction errors; however, consistently positive-going responses for punishment indicated that this prediction error signal may be unsigned. A model that puts the same positive weight on both aversive and appetitive outcomes tracks the expectation of a reinforcement irrespective of its valence; therefore, it would predict identical responses for reward- and punishment-predicting cues if omission rate is constant. However, BFCNs clearly preferred reward-predicting stimuli, suggesting differential representation of rewards and punishments. Therefore, we implemented a RL model and fitted parameters capturing differential weighing based on valence. We found that this model reliably predicted average BFCN responses to cues, rewards, and punishments. It also reproduced larger BFCN responses to cues that foreshadowed rewards with high probability, as well as to surprising, as compared to expected rewards.

What may be the function of this fast, unsigned prediction error signal? The cholinergic system has long been known to strongly influence cortical plasticity (Hasselmo and Sarter, 2011; Leão et al., 2012; Lin et al., 2015; Palacios-Filardo and Mellor, 2019; Gasselin et al., 2021; Gombkoto et al., 2021). A line of studies has demonstrated that pairing auditory stimuli with cholinergic stimulation reorganizes cortical sensory representations, known by the term ‘receptive field plasticity’ (Kilgard and Merzenich, 1998; Froemke et al., 2007). Furthermore, recent studies showed that cholinergic inputs may even endow primary sensory cortices with non-sensory representations not expected previously (Shuler and Bear, 2006; Chubykin et al., 2013). In particular, Liu et al. showed that optogenetic activation of cholinergic fibers in the visual cortex entrained neural responses that mimicked behaviorally-conditioned reward timing activity (Liu et al., 2015). It was also demonstrated that the cholinergic system exerts a rapid, fine-balanced control over plasticity at millisecond time scales, stressing the importance of timing even for neuromodulatory systems (Berg, 2011; Gu and Yakel, 2011; Urban-Ciecko et al., 2018). This effect on plasticity might have a fundamental impact on associative learning at the behavioral level (Pinto et al., 2013), also suggested by recent advances in the fear learning field (Letzkus et al., 2011; Jiang et al., 2016; Guo et al., 2019; Crouse et al., 2020).

Indeed, we found that the best fit model parameters were correlated with the difference in the animals’ anticipatory lick rate indicating learning performance. Moreover, cholinergic responses to reward-predicting cues predicted behavioral responses and reaction time, fitting in a more general scheme of basal forebrain control over response speed to motivationally salient stimuli (Lin and Nicolelis, 2008; Avila and Lin, 2014; Lin et al., 2015). Therefore, we propose that a rapid acetylcholine-mediated cortical activation, scaled by unsigned outcome prediction error, tunes synaptic plasticity in the service of behavioral learning. This idea is supported by strong theories that associated unsigned prediction errors and cholinergic activity with learning and memory (Rouhani et al., 2018; Zannone et al., 2018; Rouhani and Niv, 2021). Nevertheless, the functions of cholinergic effects probably go beyond learning, and BFCNs may control many aspects of behavior including arousal or alertness (Buzsaki et al., 1988; Richardson and DeLong, 1991; Metherate et al., 1992; Lin and Nicolelis, 2008; Zhang et al., 2011; Eggermann et al., 2014; Lin et al., 2015; Nelson and Mooney, 2016; Teles-Grilo Ruivo et al., 2017), attention (Disney et al., 2007; Parikh et al., 2007; Harris and Thiele, 2011; Hasselmo and Sarter, 2011; Pinto et al., 2013) and vigilance (Duque et al., 2000; Lee et al., 2005; Xu et al., 2015).

The activity of cholinergic neurons share strong similarities with dopaminergic neurons in response to reward and reward-predicting cues (Schultz et al., 1997; Cohen et al., 2012; Schultz, 2015; Lak et al., 2016). Reward-predicting cues evoke an increase in firing rate, which is stronger for more likely rewards. Reward itself also elicits cholinergic firing, but less so if the reward is more expected. However, cholinergic neurons differ from dopaminergic neurons in their response to punishment. While dopaminergic neurons can respond to aversive stimuli with either increased or decreased firing (Matsumoto and Hikosaka, 2009; Menegas et al., 2018; Tsutsui-Kimura et al., 2020), cholinergic neurons consistently respond with a fast, precisely timed response to air puffs. Therefore, the positive-going response of BFCNs irrespective of valence, sensitive to outcome probabilities for cues and rewards, suggests that compared to the reward prediction error signal dopaminergic neurons encode, BFCNs represent an unsigned outcome prediction error. Nevertheless, a valence-weighed unsigned prediction error hypothesis predicts stronger response to unexpected than to expected punishment; the larger the weight of the punishment, the stronger the difference. We did not find a significant difference, which could be due to low weight of punishment that decreased statistical power, or a theoretical deviation from a full-fledged outcome prediction error. Since in our design conditioned stimuli could be followed by either reward or punishment, punishment expectation signals could not be tested in isolation.

Additionally, an unsigned prediction error signal predicts a firing rate increase after omitted reward. (Note that unsigned prediction error variables only take non-negative values, and thus all unexpected changes in state value result in increased values due the absolute value operator.) However, given the phasic nature of cholinergic reinforcement responses comprising often very few (sometimes only one) but very precisely timed action potentials (Hangya et al., 2015), it is expected that an omission response, where there is no sensory stimulus to align to, is very hard to detect in single neurons. Alternatively, the cholinergic system may be sensitive to external sensory stimuli but not to absence of an expected stimulus. A recent study demonstrated topographic variations in cholinergic responses to salient events (Robert et al., 2021), which could also contribute to these ambiguities. Importantly, bursting basal forebrain neurons with similar coding properties have been uncovered in primates (Zhang et al., 2019), suggesting that at least part of those neurons might be cholinergic.

Cholinergic neurons appeared to respond faster than dopaminergic neurons; however, response timing may depend on seemingly subtle details of the behavioral paradigm. Altogether, BFCNs appear to provide a faster but less specific response to salient stimuli, which is likely broadcasted to large cortical areas innervated by cholinergic fibers (Zaborszky et al., 2013; Gielow and Zaborszky, 2017). In contrast, calculations related to value that are represented in the dopaminergic system may require more processing time and result in somewhat delayed, albeit more specific representations. Nevertheless, direct comparisons of cholinergic and dopaminergic neurons in the same experiment will be necessary to reveal the differential functions of these major neuromodulatory systems.

## Materials and methods

### Animals

Adult (over two months old) male ChAT-Cre mice (The Jackson Laboratory, RRID: IMSR_JAX:006410) were used (n = 11). All experiments were approved by the Animal Care and Use Committee of the Institute of Experimental Medicine and the Committee for Scientific Ethics of Animal Research of the National Food Chain Safety Office (PE/EA/675-4/2016; PE/EA/1212-5/2017; PE/EA/864-7/2019) and were performed according to the guidelines of the institutional ethical code and the Hungarian Act of Animal Care and Experimentation (1998; XXVIII, section 243/1998, renewed in 40/2013) in accordance with the European Directive 86/609/CEE and modified according to the Directive 2010/63/EU.

### Tetrode implantation surgery

Mice were implanted using standard stereotaxic surgery techniques with miniaturized microdrives housing 8 tetrodes and an optic fiber, constructed as described previously (Kvitsiani et al., 2013; Pi et al., 2013; Hangya et al., 2015). Briefly, mice were anesthetized by a mixture of ketamine and xylazine (83 and 17 mg/kg, respectively, dissolved in 0.9% saline). The skin was shaved and disinfected with Betadine, subcutaneous tissues were infused with Lidocaine, eyes were protected with eye ointment (Laboratories Thea) and mice were placed in a stereotaxic frame (Kopf Instruments). The skull was cleaned, and a craniotomy was drilled above the horizontal diagonal band of Broca (HDB, antero-posterior 0.75 mm, lateral 0.60 mm; n = 4) or the medial septum (MS, antero-posterior 0.90 mm, lateral 0.90 mm, 10 degrees lateral angle; n = 1). Virus injection (AAV2/5.EF1a.Dio.hChR2(H134R)-eYFP.WPRE.hGH; HDB, dorso-ventral 5.00 and 4.70 mm, 300 nl at each depth; MS, dorsoventral 3.95, 4.45 and 5.25 mm, 200 nl at each depth) and drive implantation was performed according to standard techniques as described previously (Kvitsiani et al., 2013; Hangya et al., 2015). Ground and reference electrodes were implanted to the bilateral parietal cortex. Mice received analgesics (Buprenorphine, 0.1 mg/kg), local antibiotics (Neomycin) and were allowed 10 days of recovery before starting behavioral training.

### Optical fiber implantation surgery

Mice (n = 7) were implanted using standard stereotactic surgery techniques described in the previous section. Following virus injection (AAV2/9.CAG.Flex.GCAMP6.WPRE.SV40; HDB, 300 nl each side, antero-posterior 0.75 mm, lateral 0.60 mm; dorso-ventral 5.00 and 4.7 mm), 400 µm core diameter optic fibers with ceramic ferrules were implanted bilaterally (HDB, antero-posterior 0.75 mm, lateral 0.60 mm; 0 and 20 degrees lateral angle on the two sides). Optical fiber implantation was performed similarly to tetrode drive implantation. Mice received analgesics (Buprenorphine, 0.1 mg/kg), local antibiotics (Neomycin) and were allowed 10 days of recovery before starting behavioral training.

### Behavioral training

Mice were trained on a head-fixed probabilistic auditory Pavlovian conditioning task (Hegedüs et al., 2021a) in a custom-built behavioral setup that allowed millisecond precision of stimulus and reinforcement delivery (described in (Solari et al., 2018)). Mice were water restricted before training and worked for small amounts of water reward (5 µl) during conditioning. Pure tones of one second duration predicted likely reward / unlikely punishment or unlikely reward / likely punishment based on their pitch (12 kHz tone predicted 80% water reward, 10% air-puff punishment, 10% omission; 4 kHz tone predicted 25% water reward, 65% air-puff punishment, 10% omission in n = 6 mice; opposite cue contingencies were used in n = 5 mice, see Fig. S1 and S3; 50-50% of the two cue tones were mixed randomly). The animal was free to lick a waterspout after tone onset and individual licks were detected by the animal’s tongue breaking an infrared photobeam. After an additional 200-400 ms post-stimulus delay, the animal received water reward, air-puff punishment or omission, pseudorandomized according to the above contingencies. The next trial started after the animal stopped licking for at least 1.5 s. The stimulus was preceded by a 1-4 s foreperiod according to a truncated exponential distribution, in order to prevent temporal expectation of stimulus delivery. If the mouse licked in the foreperiod, the trial was restarted. We used the open source Bpod behavioral control system (Sanworks LLC, US) for operating the task. Behavioral performance of the task did not depend on the identity (frequencies) of the conditioned stimuli (Fig. S1).

The aversive quality of air-puffs depends on the exact experimental settings. We applied 200 ms long puffs at 15 psi pressure (within the range of parameters used for eyeblink conditioning (Najafi et al., 2014)). We demonstrated that mice consistently choose water without air-puff over water combined with air-puff, showing that air-puffs are aversive under these circumstances (see Fig. 2C-D in (Hangya et al., 2015)). We also demonstrated that water and air-puff are accompanied by different auditory signals in our setup, thus making sensory response generalization unlikely to explain BFCN responses (see Fig. S2A in (Hegedüs et al., 2021a)).

### Fiber photometry imaging

Bilateral fluorescent calcium imaging was performed using a dual fiber photometry setup (Doric Neuroscience) and visualized during training sessions using Doric Studio Software. Two LED light sources (465 nm, 405 nm) were channeled in fluorescent Mini Cubes (iFMC4, Doric Neuroscience). Light was amplitude-modulated by the command voltage of the two-channel LED driver (LEDD_2, Doric Neuroscience, the 465 nm wavelength light was modulated at 208 Hz and 405 nm wavelength was modulated at 572 Hz). Light was channeled into 400 µm diameter patch cord fibers and was connected to optical fiber implants during training sessions. The same optical fibers were used to collect the bilateral emitted fluorescence signal, which were detected with 500-550 nm fluorescent detectors integrated in the Mini Cubes. Emitted signals were sampled at 12 kHz, decoded in silico and saved in a *.csv format.

### Chronic extracellular recording

We used the open ephys data acquisition system (Siegle et al., 2017) for spike data collection. A 32-channels Intan headstage (RHD2132) was connected to the Omnetics connector on the custom-built microdrive. Data was transferred through digital SPI cables (Intan) to the Open Ephys board and saved by the Open Ephys software, digitized at 30 kHz.

### Optogenetic tagging

The custom microdrives were equipped with a 50 μm core optic fiber (Thorlabs) that ended in an FC connector (Precision Fiber Products). This was connected with an FC-APC patch chord during recording. For optogenetic tagging, 1 ms laser pulses were delivered (473 nm, Sanctity) at 20 Hz for 2 seconds, followed by 3 seconds pause, repeated 20-30 times. Light-evoked spikes and potential artifacts were monitored online using the OPETH plugin (Széll et al., 2020) and laser power was adjusted as necessary to avoid light-induced photoelectric artifacts and population spikes that could mask individual action potentials. Significance of photoactivation was assessed during offline analyses by the SALT test based spike latency distributions after light pulses, compared to a surrogate distribution using Jensen-Shannon divergence (information radius) (Endres and Schindelin, 2003) as described previously (Kvitsiani et al., 2013). Neurons with p < 0.01 were considered light-activated, and thus cholinergic. Cholinergic neurons recorded on the same tetrode within 200 μm dorso-ventral distance were compared by waveform correlation and autocorrelogram similarity (Fraser and Schwartz, 2012; Hegedüs et al., 2021a), and similar units were counted towards the sample size only once.

### Histology

After the last behavioral session, mice were deeply anesthetized with ketamine/xylazine and we performed an electrolytic lesion to aid electrode localization (5 uA current for ∼5s on 2 leads/tetrode), Supertech, IBP-7c). Mice were perfused transcardially, starting with a 2-minute washout period with saline, followed by 4% paraformaldehyde solution for 20 minutes. After the perfusion, mice were removed from the platform and decapitated. The brain was carefully removed and postfixed overnight in 4% PFA. A block containing the full extent of the HDB was prepared and 50 µm thick sections were cut using a Leica 2100S vibratome. All attempts were made to section parallel to the canonical coronal plane to aid track reconstruction efforts. All sections that contained the electrode tracks were mounted on slides in Aquamount mounting medium. Fluorescent and dark field confocal images of the sections were taken with a Nikon C2 confocal microscope. During track reconstruction, it is important to convert the logged screw turns (20 μm for each one eights of a full turn, allowed by a 160 μm pitch custom precision screw, Easterntec, Shanghai) that were performed throughout the experiment into brain atlas coordinates with maximal possible precision. To this end, dark field and bright field images of the brain sections were morphed onto the corresponding atlas planes (Franklin and Paxinos, 2007) using Euclidean transformations only. The aligned atlas images were carried over to fluorescent images of the brain sections showing the DiI-labelled electrode tracks (red) and green fluorescent labeling (cholinergic neurons labelled by the AAV2.5-EF1a-Dio-hChR2(H134R)-eYFP.WPRE.hGh virus) in the target area. The entry points, electrode tips and lesion sites were localized with respect to the atlas coordinates maximizing the combined information of the structural (dark/bright field), DiI track and ChAT-labelling fluoromicrographs. Recording location of each section was interpolated based on the above coordinates, using the screw turn logs and the measured protruding length of the tetrodes after the experiments (also described in (Hangya et al., 2015)). If the track spanned multiple sections, special care was taken to precisely reconstruct the part of the track where the recordings took place within the target area. This procedure minimizes the localization errors that may arise from differences between the recorded and the reference brain coordinates and eliminates the effect of tissue distortions caused by the fixation process. Only those recordings that were convincingly localized to the basal forebrain were analyzed in this study.

### Data analysis

Data processing and analysis was carried out in Matlab R2016a (Mathworks, Natick).

Fiber photometry signals were preprocessed according to Lerner et al. (2015). In case of bilateral data acquisition, fiber photometry signal of the side with the better signal-to-noise ratio was chosen for further analysis. Briefly, the fluorescence signals were filtered below 20 Hz using a low-pass Butterworth digital filter to remove high frequency noise. To calculate dff, a least-squares linear fit was applied to the isosbestic 405 nm signal to align its baseline intensity to that of the calcium-dependent 465 nm (f465) signal. The fitted 405 nm signal (f_405,fitted_) was used to normalize the 465 nm signal as follows:

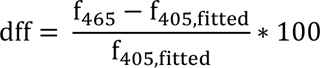

to remove the effect of motion and autofluorescence. Slow decay of the baseline activity was filtered out with an 0.2 Hz high pass Butterworth digital filter. Finally, the dff signal was triggered on cue and feedback times, Z-scored by the mean and standard deviation of a baseline window (1s before cue onset) and averaged across trials.

Tetrode recording channels were digitally referenced to a common average reference, filtered between 700-7000 Hz with Butterworth zero-phase filter and spikes were detected using a 750 µs censoring period. Spike sorting was carried out in MClust 3.5 software (A .D. Redish). Autocorrelations were inspected for refractory period violations and putative units with insufficient refractory period were not included in the data set. Cluster separation was measured using Isolation Distance and L-ratio calculated on the basis of two features, the full spike amplitude and the first principle component of the waveform (Schmitzer-Torbert et al., 2005). Putative single neurons exceeding Isolation Distance (ID) of 20 and below L-ratio of 0.15 were automatically included (n = 20 cholinergic neurons and n = 452 untagged neurons recorded in the same sessions). Additionally, spike sorting was aided by the information provided by light-evoked spike shapes (Pi et al., 2013) in n = 5 cholinergic neurons, resulting in a data set of n = 25 optogenetically identified cholinergic neurons (L-ratio of cholinergic neurons, 0.0511 ± 0.0133, median ± SE; ID of cholinergic neurons, 30.0809 ± 6.4121, median ± SE; n = 17, 5, 2, 1 cells in the four mice). Spike shape correlations between spontaneous and light-induced spikes were calculated for all cholinergic neurons. Correlation coefficient exceeded R = 0.85 in all and R = 0.9 in 22/25 optotagged neurons (0.98 ± 0.01, median ± SE; range, 0.87 - 1.0), confirming cholinergic identity (Cohen et al., 2012).

We did not find any systematic differences in our analyses based on anatomical location; thus, we analyzed the 25 neurons as one dataset. First, we calculated event-aligned raster plots and peri-event time histograms (PETHs) for all neurons. To calculate average PETHs, neuronal responses were triggered on cue and feedback times, Z-scored by the mean and standard deviation of a baseline window (1s before cue onset) and averaged across trials. Response latency and jitter to optogenetic stimulation and behaviorally relevant events were determined based on activation peaks in the peri-event time histograms as described previously (Hangya et al., 2015). Behavioral performance was tested by comparing the anticipatory lick rate after reward and punishment predicting stimuli in a 1.2 s time window after stimulus onset. Reaction time to reward was determined as the latency of the first lick after stimulus presentation.

We would like to note that an initial analysis of a part of this data set was presented in a bioRxiv preprint (https://www.biorxiv.org/content/10.1101/2020.02.17.953141v1).

### Model fitting

Firing rates of cholinergic neurons were calculated in 500 ms response windows after cue presentation and 200 ms response windows after reinforcement presentation, to include the full firing response based on the observed time course of cholinergic activation (Fig. 4). Firing rates were fitted by the following modified temporal difference reinforcement learning model of cholinergic activity *(C)*.

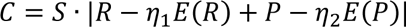

In this equation, 𝑅 − 𝜂_1_𝐸(𝑅) stands for reward prediction error (RPE). RPE classically takes the formula of 𝑅 − 𝐸(𝑅), where 𝐸(𝑅) is expected, and 𝑅 is actual amount of reward at a given time point. This was modified by the *η_1_* parameter, allowing potential differences in sensitivity to reward expectation across animals, sessions and neurons. Similarly, 𝑃 − 𝜂_2_𝐸(𝑃) represents the difference of expected and encountered punishment, referred to as ‘punishment prediction error’ hereafter. The two terms sum up to a full outcome prediction error, rendered ‘unsigned’ by the absolute value operator. The scaling factor *S* accounts for differences in baseline firing rate of cholinergic neurons. The temporal discounting factor inherent to TDRL models was omitted from the equation, as it leads to only negligible firing rate differences within the few seconds of time that spans a behavioral trial. We note that another way of incorporating differential responsiveness to reward and punishment expectations would be by adding classical learning rates in the form of

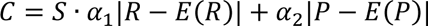

We fitted this alternative model as well; however, this model failed to capture the relative ratios of cue and outcome responses of cholinergic neurons and thus resulted in worse fits than the model presented in Fig. 5.

The model was evaluated for the time of the cue, reward and punishment presentations. At the time of cues, *R = P = 0*, therefore the model takes the form of

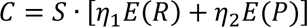

Since no omission responses were observed in cholinergic recordings (Fig. S7), we dropped the negative expectation term of the omitted reinforcer at the time of reinforcement (e.g. omitted reward response at the time of punishment), leading to

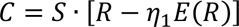

at reward (*P = 0*) and

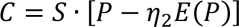

at punishment (*R = 0*) delivery. Nevertheless, keeping the omission responses in the model resulted in fits that were statistically indifferentiable (p = 0.86, Wilcoxon signed-rank test of model errors), suggesting that our data were not sufficient to differentiate between RL models with or without omission responses. The *E(R)* and *E(P)* expectation terms were set according to the task contingencies (*E(R)* = 0.8 or 0.25 and *E(P)* = 0.1 or 0.65 for the likely reward and likely punishment cues, respectively). As a control model, we ran the same fitting process after these contingencies for reward and punishment expectations were swapped (*E(R)* = 0.25 or 0.8 and *E(P)* = 0.65 or 0.1 for the likely reward and likely punishment cues, respectively). Fitting error was estimated by the maximum likelihood method and minimized by using the fminsearch built-in Matlab function employing the Nelder-Mead simplex algorithm. Models were statistically compared by Wilcoxon signed-rank test on the maximum likelihoods. Note that the compared models had equal complexity and number of parameters; therefore, a punishment term for free parameters was not required. Correlation between model parameters and anticipatory lick rate difference was calculated using the built-in robust regression algorithm of Matlab. Confidence intervals were derived using the polypredci.m function (Star Strider, https://www.mathworks.com/matlabcentral/fileexchange/57630-polypredci, MATLAB Central File Exchange, retrieved December 30, 2020).

Spike trains were simulated as Poisson-processes matched to each recorded cholinergic neuron in frequency (n = 25). Cholinergic responses to cue, reward and punishment were simulated as additional spikes drawn from a Gaussian distribution with fixed latency after the events. The number of ‘evoked spikes’ was based on the best-fit RL model corresponding to each neuron. Peri-event time histograms were generated from simulated spike trains the same way as applied for real data.

### Statistics

We estimated the sample size before conducting the study based on previous publications, mostly (Hangya et al., 2015), as reported in the Results. Firing rates and other variables were compared across conditions using non-parametric tests, as normality of the underlying distributions could not be determined. Two-sided Wilcoxon signed-rank test was applied for paired, and two-sided Mann-Whitney U-test was applied for non-paired samples. Correlations were estimated by the Pearson’s correlation coefficient, and their significance were judged by using a standard linear regression approach (one-sided F-test, in accordance with the asymmetric null hypothesis of linear regression). Model fits were compared by negative log likelihood. Since the models compared had equal number of parameters, this is mathematically equivalent with model selection approaches using information criteria (e.g. Akaike and Bayesian Information Criterion). Peri-event time histograms show mean ± SE. Box-whisker plots show median, interquartile range and non-outlier range, with all data point overlaid.

## Author contributions

PH and BH conceived the study; PH and KS performed the recording experiments; PH and SMB conducted the fiber photometry experiments; PH, BH and BK analyzed the data; PH prepared the figures, BH and PH wrote the manuscript with inputs from KS.

## Data availability

Electrophysiology and fiber photometry data have been deposited to a Dryad repository at https://doi.org/10.5061/dryad.p5hqbzkrv. Please kindly note that while this link is not live yet, the dataset is accessible through this temporary link for review: https://datadryad.org/stash/share/O0Zmp7WR-ksh8UofxgcF40hVIZMnXChOaDO0nMRTm6Y.

## Code availability

MATLAB codes generated for this study are available at https://github.com/hangyabalazs/cholinergic_Pavlovian_analysis.

## Acknowledgements

We thank Katalin Lengyel for technical assistance in anatomical methods, Annal Velencei and Victoria Lyakhova for assistance in behavioral training and Dr. Janos Szabadics for helpful comments on the manuscript. This work was supported by the ‘Lendület’ Program of the Hungarian Academy of Sciences (LP2015-2/2015), the Artificial Intelligence National Laboratory Programme of the Ministry for Innovation and Technology, NKFIH KH125294, NKFIH K135561, SPIRITS 2020 of Kyoto University and the European Research Council Starting Grant no. 715043 to BH; the ÚNKP-21-3 New National Excellence Program of the Ministry for Innovation and Technology to PH; ÚNKP-19-3, ÚNKP-20-3 and ÚNKP 21-3 New National Excellence Program of the Ministry for Innovation and Technology from the source of the National Research, Development and Innovation Fund to BK and the Generalitat Valenciana Postdoctoral Fellowship Program (APOSTD/2019/003) to SMB. We acknowledge the help of the Nikon Center of Excellence at the Institute of Experimental Medicine (IEM), Nikon Europe, Nikon Austria and Auro-Science Consulting for kindly providing microscopy support and the supportive help of the Central Virus Laboratory of IEM. We thank Mackenzie Mathis for open access science art at SciDraw (accessible at https://doi.org/10.5281/zenodo.3925907).

## Supplemental information

**Figure S1.**
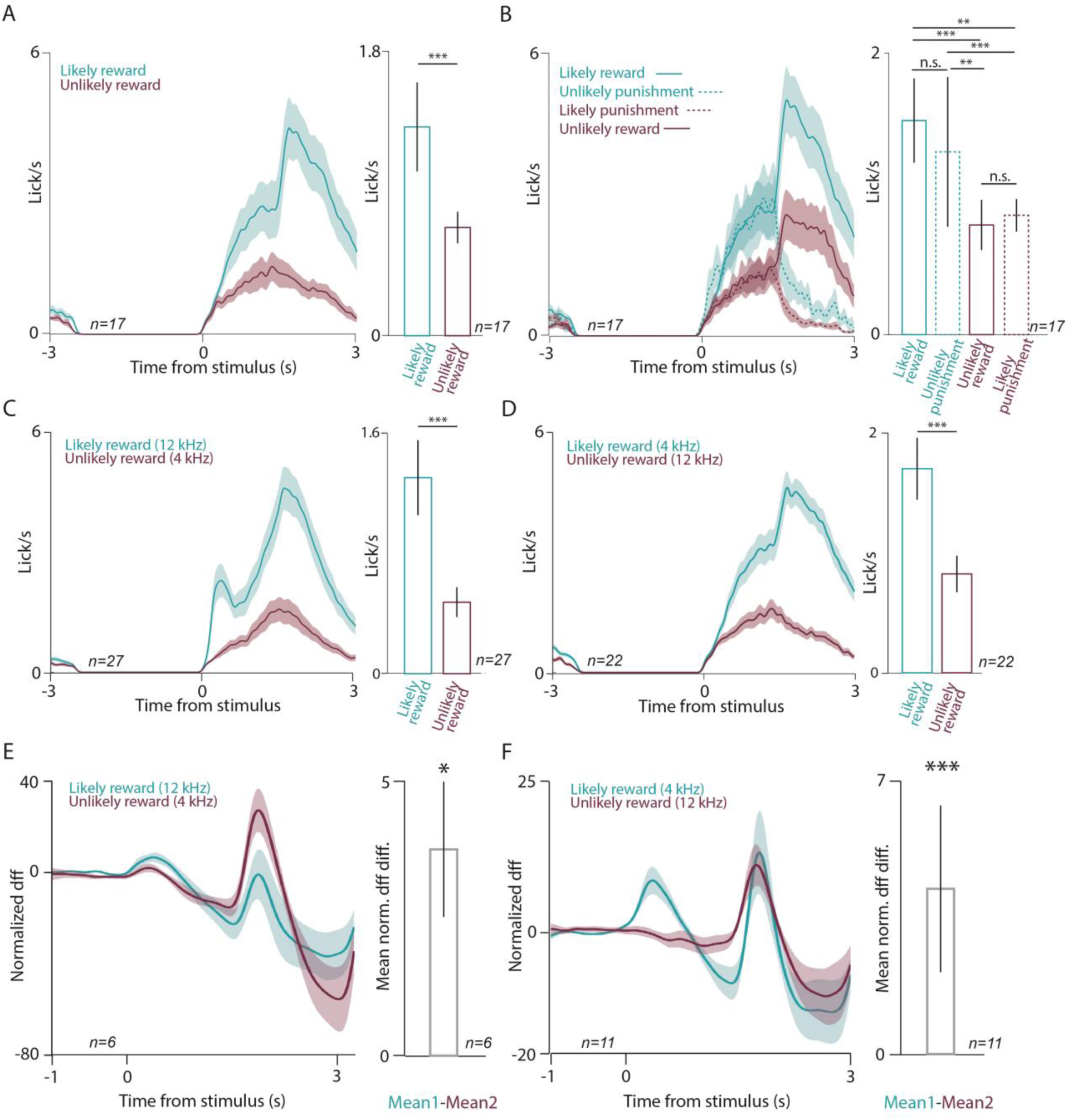
Behavioral performance during bulk calcium imaging of cholinergic neurons. (A) Left, average PETHs of lick responses of all sessions during fiber photometry imaging (n = 17 sessions). Right, statistical comparison of anticipatory lick rates in the RW in likely reward and unlikely reward trials (n = 17 sessions, median ± SE of median, ***, p < 0.001, p = 0.00042, Wilcoxon signed-rank test). (B) Left, average PETHs of lick responses of all sessions of all animals (n = 17 sessions), partitioned based on the four possible outcomes: expected or surprising reward, expected or surprising punishment. Right, statistical comparison of anticipatory lick rates in the RW with respect to the four possible outcomes (n = 17 sessions, median ± SE of median, **, p < 0.01, ***, p < 0.001, n.s., p > 0.05; from top to bottom, p = 0.00118, p = 0.00085, p = 0.00042, p = 0.7226, p = 0.0012, p = 0.9811, Wilcoxon signed-rank test). (C) Left, average PETHs of lick responses in sessions where likely reward (unlikely punishment) was predicted by a 12 kHz tone and unlikely reward (likely punishment) was predicted by a 4 kHz tone (n = 27 sessions, all sessions with full task contingencies were included). Right, statistical comparison of anticipatory lick rates in the RW in likely reward and unlikely reward trials (n = 27 sessions, median ± SE of median, ***, p < 0.001, p = 2.91 x 10^-5^, Wilcoxon signed-rank test). (D) The same as C, but with opposite contingencies (n = 22 sessions, all sessions with full task contingencies were included; median ± SE of median, ***, p < 0.001, p = 0.00033, Wilcoxon signed-rank test). (E) Left, average PETH of Z-scored dff aligned to outcome-predicting conditioned stimuli, where likely reward (unlikely punishment) was predicted by a 12 kHz tone and unlikely reward (likely punishment) was predicted by a 4 kHz tone (n = 6 sessions). Right, bar graph of average normalized difference in dff after likely reward and unlikely reward cues. Median ± SE of median, *, p < 0.05, p = 0.03125, Wilcoxon signed-rank test. (F) The same as E, but with opposite contingencies (n = 11 sessions, median ± SE of median, ***, p < 0.001, p = 0.00098, Wilcoxon signed-rank test).

**Figure S2.**
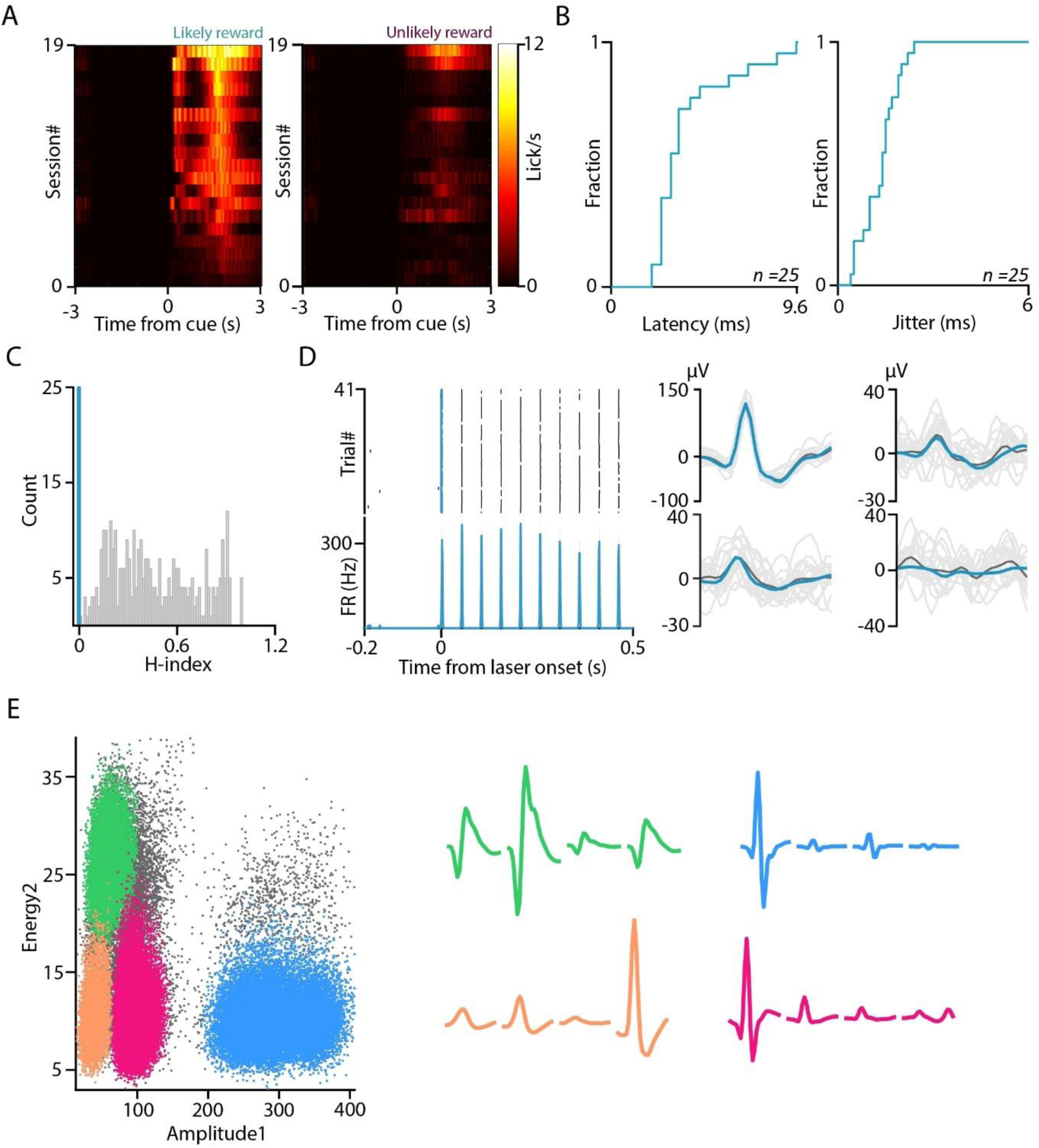
Optogenetic tagging of cholinergic neurons. (A) Color-coded PETHs of lick responses of all sessions in which cholinergic neurons were recorded (n = 19 sessions; black, no licks; white, maximal lick response). (B) Left, cumulative histogram of the peak response latency of BFCNS after optogenetic stimulation. Right, cumulative histogram of the jitter of cholinergic spike responses after optogenetic stimulation. (C) Distribution of the significance values of the SALT statistical test (H-index) for all recorded neurons (blue, p < 0.01, tagged cholinergic neurons; grey, p > 0.01, untagged neurons). (D) Left, example spike raster (top) and PETH (bottom) of an optogenetically tagged BFCN responding to 20 Hz blue laser light stimulation. Right, average spike waveform of the same BFCN on the four tetrode channels (blue, average light-evoked spikes; black, average spontaneous spikes; grey, all spikes). (E) Left, example of spike clusters plotted in feature space from a recording session. Right, average spike waveform of the recorded neurons on each tetrode channel.

**Figure S3.**
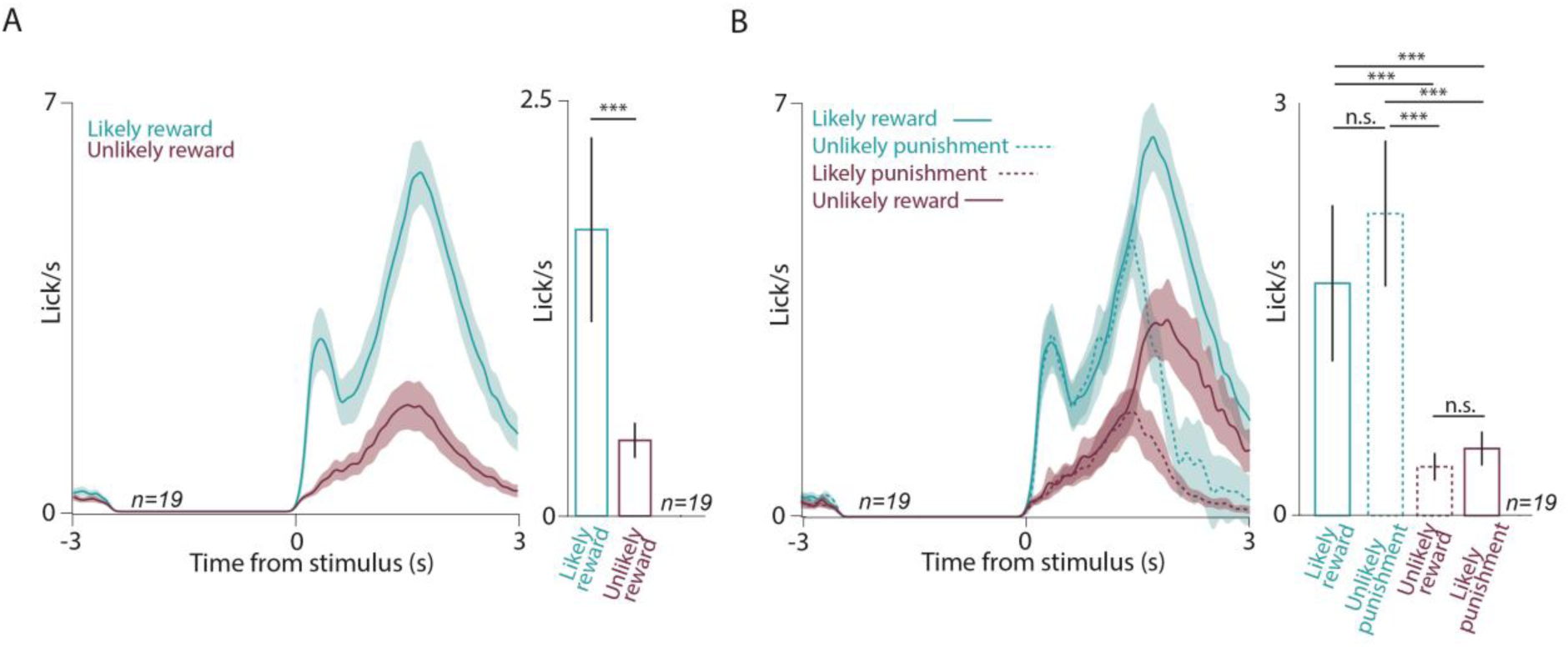
Behavioral performance during tetrode recordings. (A) Left, average PETHs of lick responses of all sessions of all animals (n = 19 sessions). Right, statistical comparison of anticipatory lick rates in the response window (RW) in likely reward and unlikely reward trials (median ± SE of median, n = 19 sessions; ***, p < 0.001, p = 0.00034, Wilcoxon signed-rank test). (B) Left, average PETHs of lick responses of all sessions of all animals (n = 19 sessions), partitioned based on the four possible outcomes: expected or surprising reward, expected or surprising punishment. Right, statistical comparison of anticipatory lick rates in the RW, with respect to the four possible outcomes (median ± SE of median, n = 19 sessions; ***, p < 0.001, n.s., p > 0.05; from top to bottom, p = 0.00034, p = 0.00084, p = 0.00021, p = 0.8721, p = 0.00025 and p = 0.9512, Wilcoxon signed-rank test).

**Fig S4.**
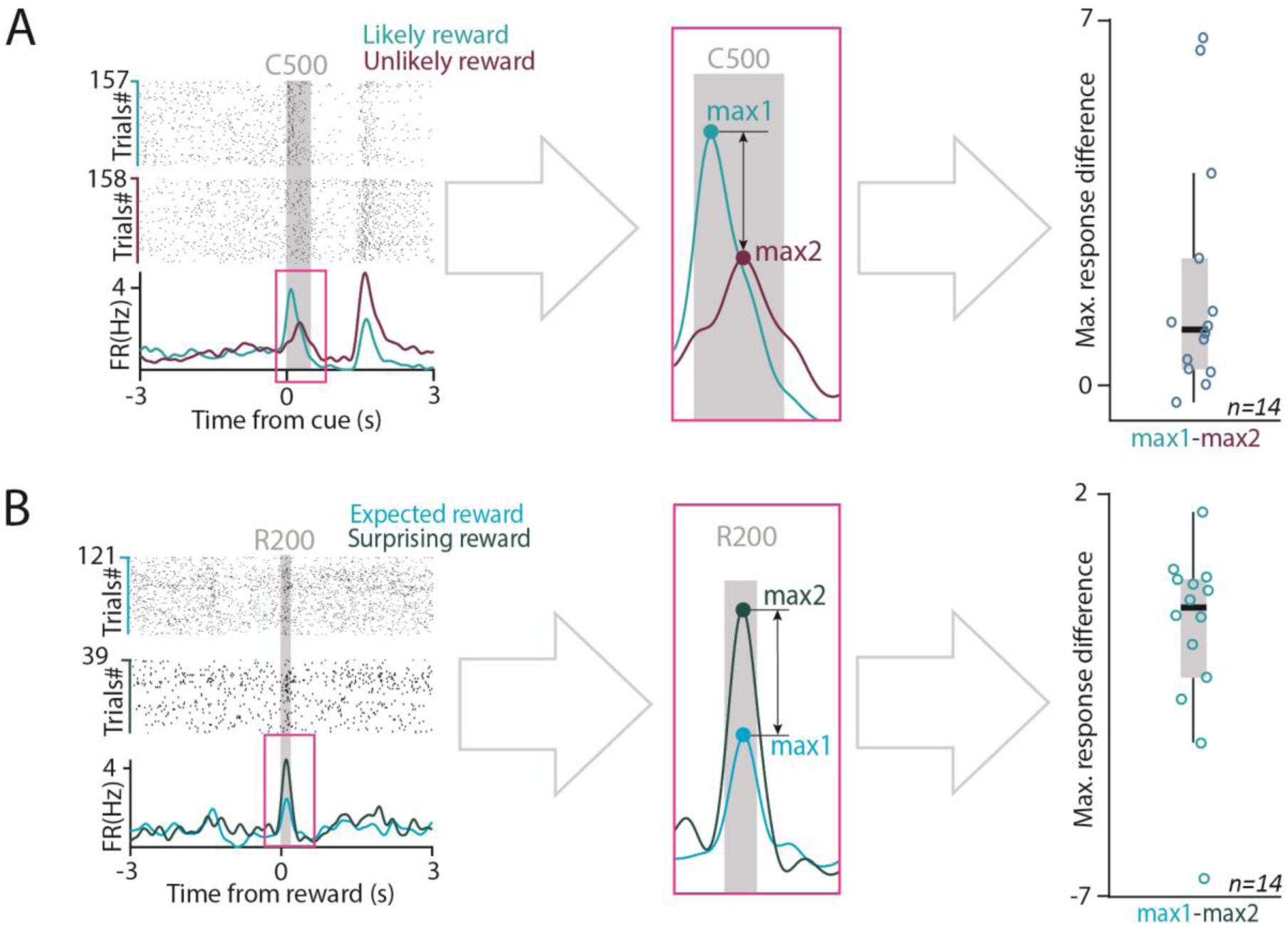
Schematic illustration of quantifying neuronal responses. (A) Schematic illustration of quantifying neuronal response to conditioned stimuli. Peak firing rate of the PETH and total spike number were calculated in a 500 ms window after cue presentation (C500), based on the range of response latencies (see Results). (B) Schematic illustration of quantifying neuronal responses to reward. Peak firing rate of the PETH and total spike number were calculated in a 200 ms window after reward delivery (R200), based on the range of response latencies (see Results). Both maximal response and average firing rate (FR) were statistically compared using Wilcoxon signed-rank test.

**Fig S5.**
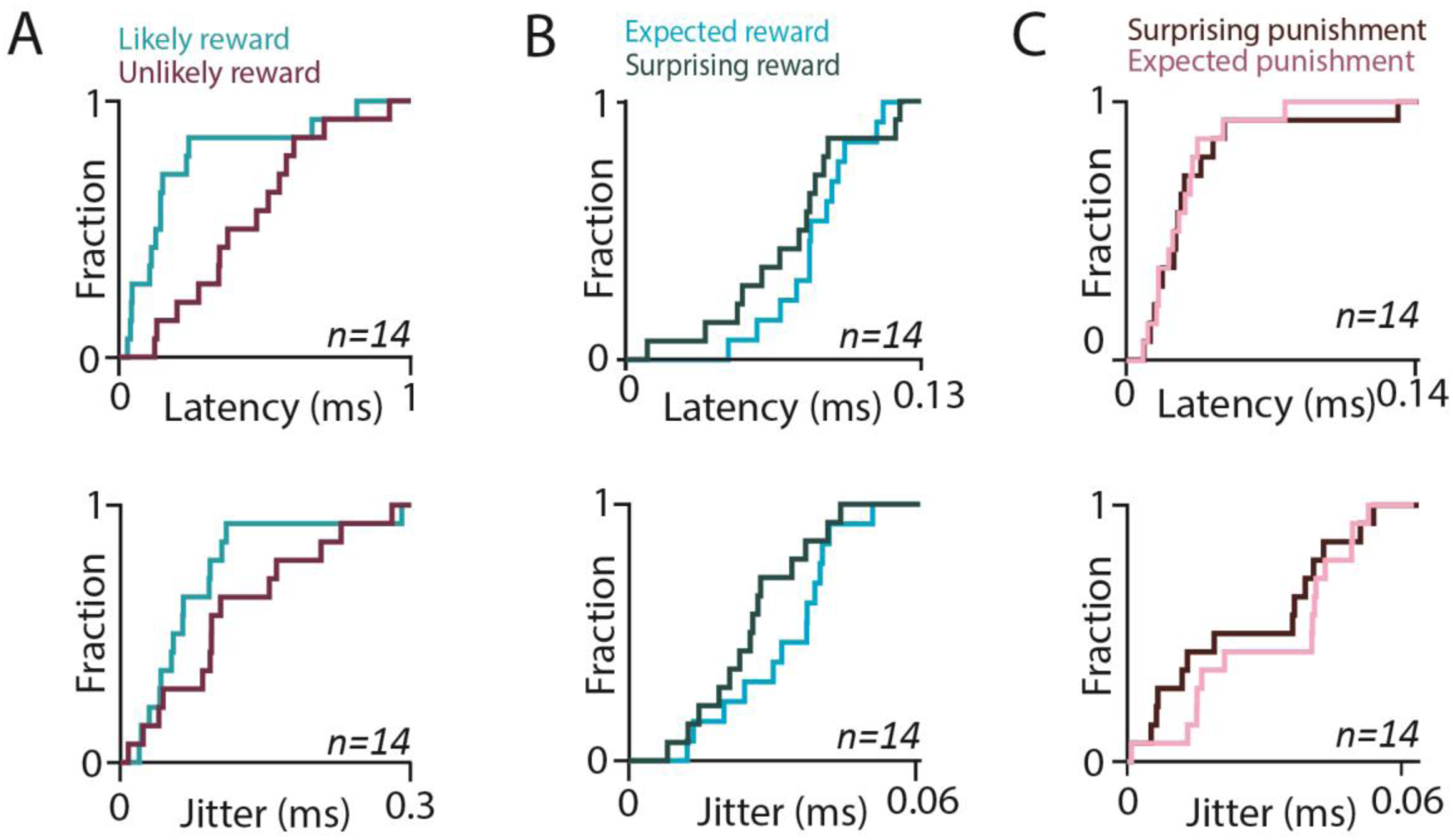
Latency and jitter of identified cholinergic neurons. (A) Top, Cumulative histogram of the peak response latencies of BFCNs (n = 14) after cue presentation. Bottom, Cumulative histogram of the jitter of cholinergic spike responses after cue presentation (n = 14 BFCNs). (B) Top, Cumulative histogram of the peak response latencies of BFCNs (n = 14) after reward. Bottom, Cumulative histogram of the jitter of cholinergic spike responses after reward (n = 14 BFCNs). (C) Top, Cumulative histogram of the peak response latencies of BFCNs (n = 14) after punishment. Bottom, Cumulative histogram of the jitter of cholinergic spike responses after punishment (n = 14 BFCNs).

**Figure S6.**
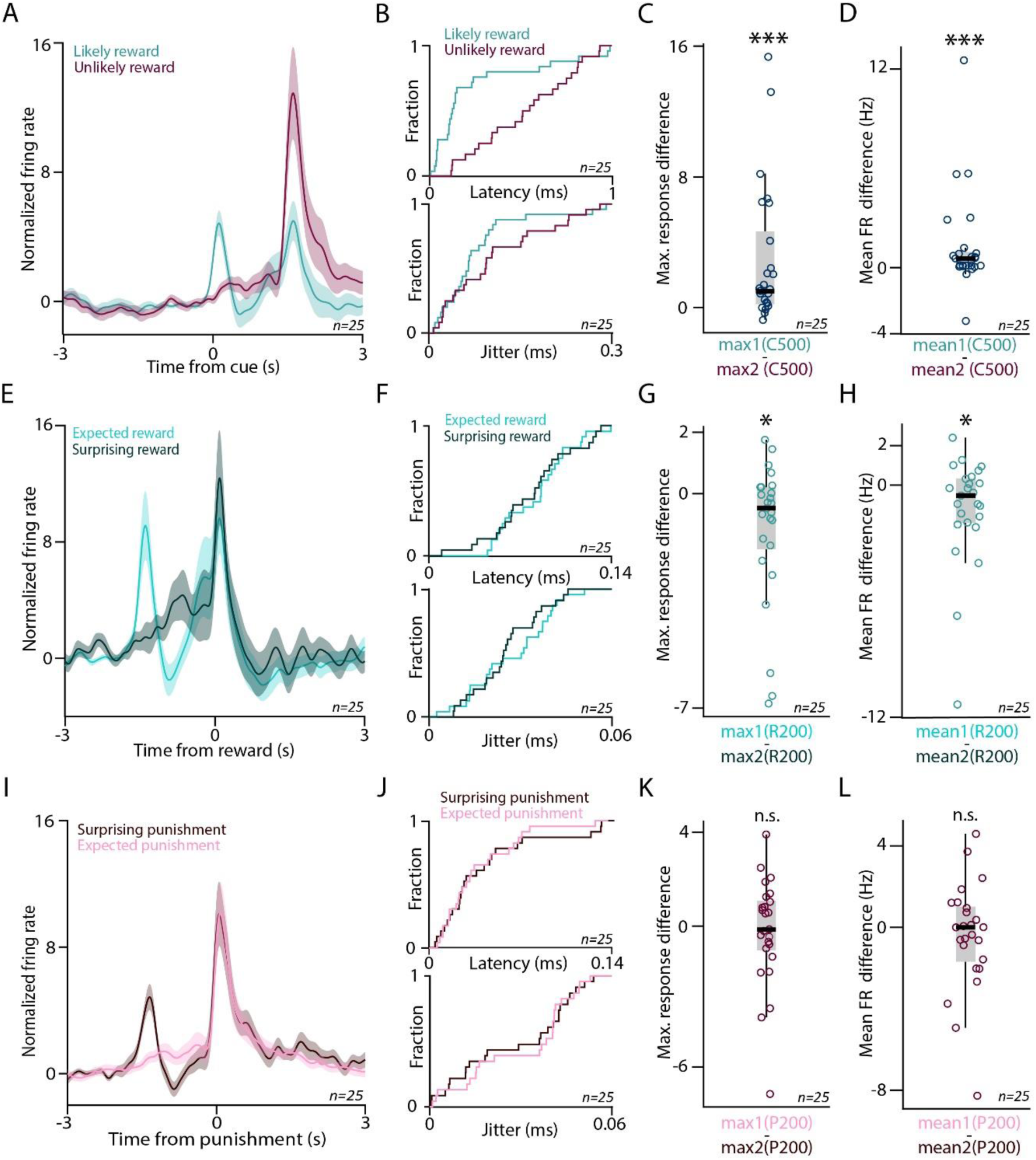
Cholinergic neurons respond more to reward-predicting cues and surprising reward. (A) Average cue-aligned PETHs of identified BFCNs (errorshade, mean ± SE; n = 25), aligned to cue onset, separately for the cues predicting likely reward (turquoise) vs. unlikely reward (purple). (B) Top, Cumulative histogram of the peak response latencies of BFCNs (n = 25) after cue presentation. Bottom, Cumulative histogram of the jitter of cholinergic spike responses after cue presentation (n = 25 BFCNs). (C) Difference in peak response after cues predicting likely reward and those predicting unlikely reward. ***, p < 0.001, p = 0.0001743, Wilcoxon signed-rank test, n = 25. (D) Difference in mean firing rate after cues predicting likely reward and those predicting unlikely reward. ***, p < 0.001, p = 0.000664, Wilcoxon signed-rank test, n = 25. (E) Average reward-aligned PETHs of identified BFCNs (errorshade, mean ± SE; n = 25), separately for rewards after the cue predicting likely reward (light turquoise, expected reward) vs. after the cue predicting unlikely reward (dark turquoise, surprising reward). (F) Top, Cumulative histogram of the peak response latencies of BFCNs (n = 25) after reward delivery. Bottom, Cumulative histogram of the jitter of cholinergic spike responses after reward delivery (n = 25 BFCNs). (G) Difference in peak response after expected and surprising rewards. *, p < 0.05, p = 0.02465, Wilcoxon signed-rank test, n = 25. (H) Difference in mean firing rate after expected and surprising rewards. *, p < 0.05, p = 0.03704, Wilcoxon signed-rank test, n = 25. (I) Average punishment-aligned PETHs of identified BFCNs (errorshade, mean ± SE; n = 25), separately for punishments after the cue predicting likely reward (dark purple, surprising punishment) vs. after the cue predicting unlikely reward (light purple, expected punishment). (J) Top, Cumulative histogram of the peak response latencies of BFCNs (n = 25) after punishment delivery. Bottom, Cumulative histogram of the jitter of cholinergic spike responses after punishment delivery (n = 25 BFCNs). (K) Difference in peak response after expected and surprising punishments. n.s., p > 0.05, p = 0.94637, Wilcoxon signed-rank test, n = 25. (L) Difference in mean firing rate after expected and surprising punishments. n.s., p > 0.05, p = 0.54851, Wilcoxon signed-rank test, n = 25.

**Figure S7.**
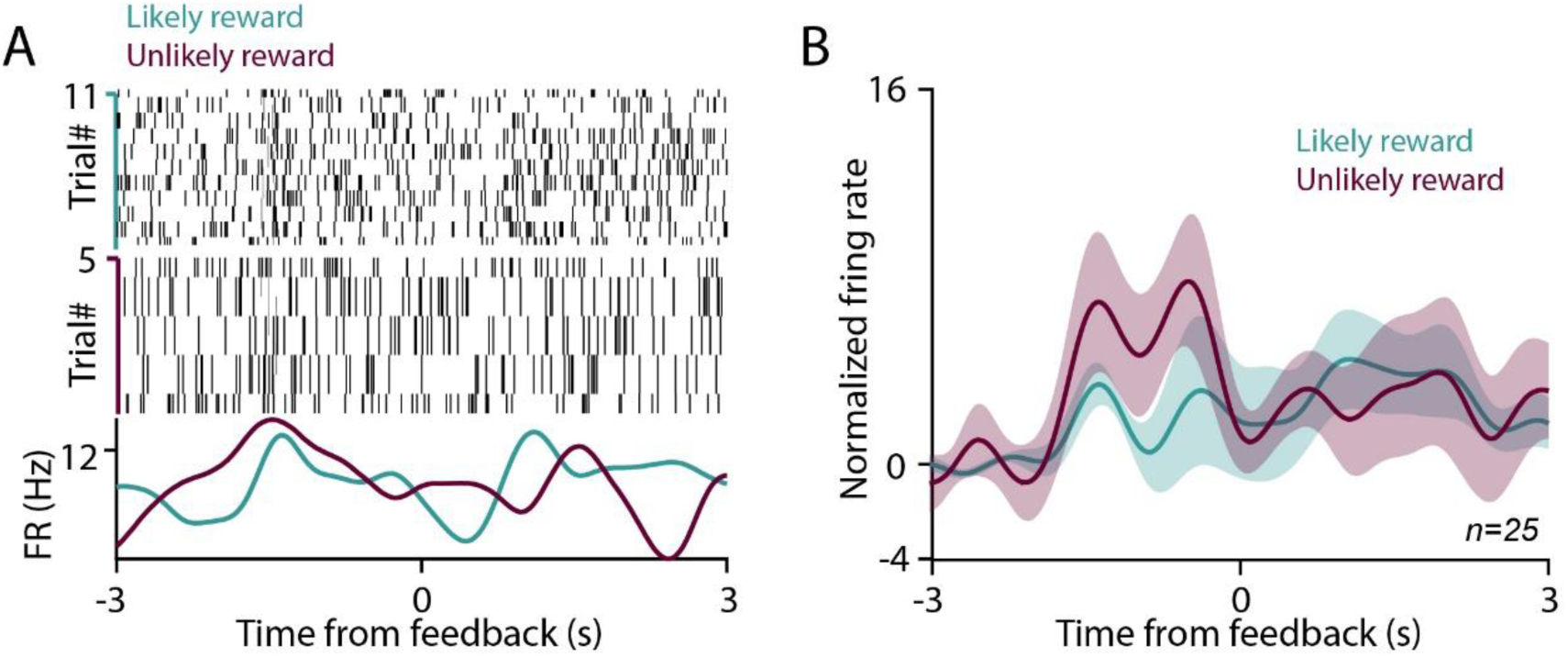
Cholinergic neurons do not respond to outcome omissions. **(A)** Example spike raster and PETH of a BFCN aligned to the expected time of omitted feedback. **(B)** Average PETH of all BFCNS (errorshade, mean ± SE; n = 25) with the same alignment.

**Figure S8.**
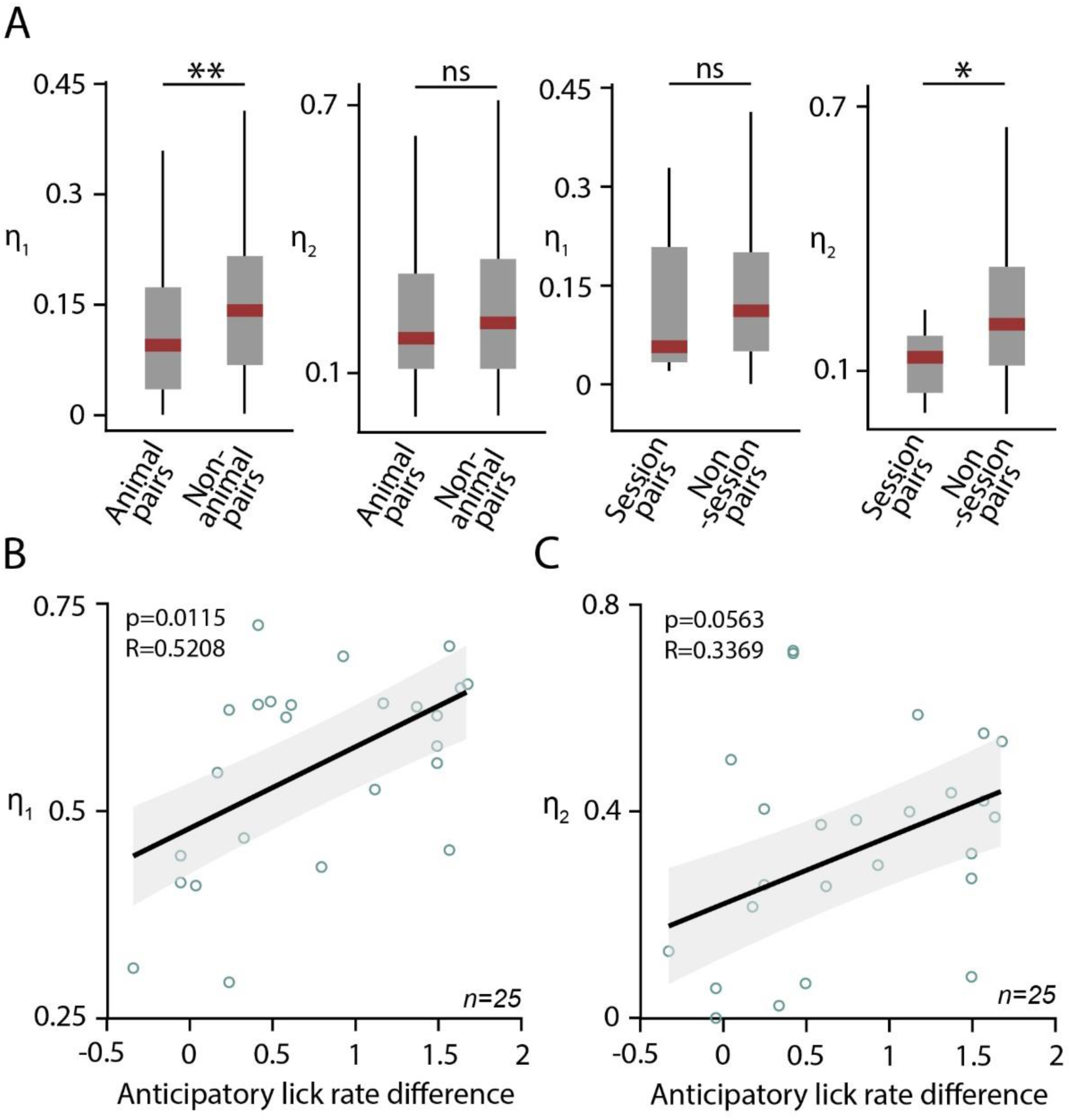
Prediction error parameters reveal behavioral correlations. **(A)** Left, difference of best-fit *η_1_* and *η_2_* parameters between BFCNs recorded in the same animal vs. in different mice. Right, difference of best-fit *η_1_* and *η_2_* parameters between BFCNs recorded in the same recording session vs. in different sessions. *, p < 0.05; **, p < 0.01, n.s., p > 0.05, Mann-Whitney U-test, n = 147 animal pairs, n = 153 non-animal pairs, n = 7 session pairs, n = 293 non-session pairs; exact p values, p = 0.00199, p = 0.2496, p = 0.5725, p = 0.04677, respectively. **(B)** Correlation between the best-fit *η_1_* parameter and the difference in anticipatory lick rate after likely reward vs. unlikely reward cues (R = 0.5208, Pearson’s correlation coefficient; p = 0.0115, linear regression, F-test). Correlation between the best-fit *η_2_* parameter and the difference in anticipatory lick rate after likely reward vs. unlikely reward cues (R = 0.3369, Pearson’s correlation coefficient; p = 0.0563, linear regression, F-test).

**Figure S9.**
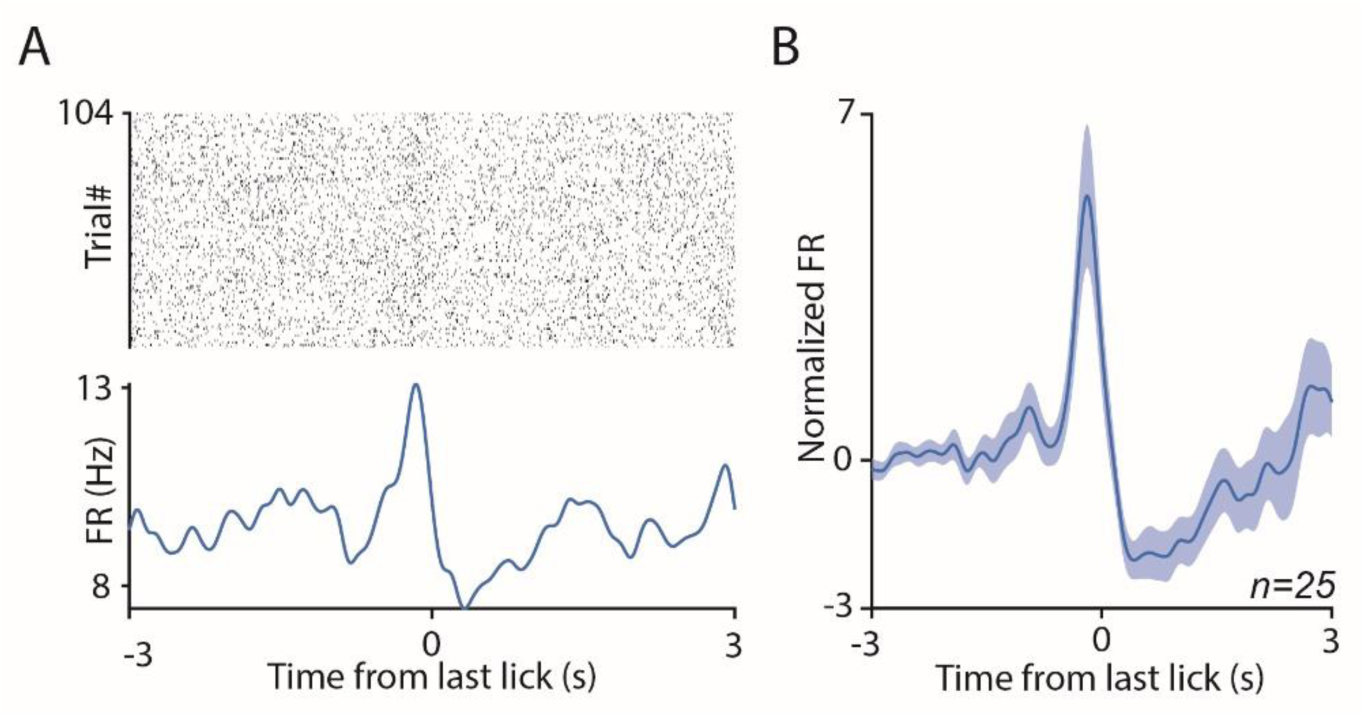
Cholinergic activity aligned to last lick before the foreperiod. (A) Raster plot (top) and PETHs (bottom) of an example cholinergic neuron aligned to the last lick before the foreperiod. (B) Average PETH of all cholinergic neurons (errorshade, mean ± SE, n = 25) aligned to the last lick before the foreperiod.
FR, firing rate.

## References

Ahrens AM, Ferguson LM, Robinson TE, Aldridge JW (2018) Dynamic encoding of incentive salience in the ventral pallidum: Dependence on the form of the reward cue. eNeuro 5:1–16.

Arendt T, Bigl V (1986) Alzheimer plaques and cortical cholinergic innervation. Neuroscience 17:277– 279.

Avila I, Lin S-C (2014) Motivational Salience Signal in the Basal Forebrain Is Coupled with Faster and More Precise Decision Speed Posner M, ed. PLoS Biol 12:e1001811.

Bailey AM, Rudisill ML, Hoof EJ, Loving ML (2003) 192 IgG-saporin lesions to the nucleus basalis magnocellularis (nBM) disrupt acquisition of learning set formation. Brain Res 969:147–159.

Berg DK (2011) Timing is everything, even for cholinergic control. Neuron 71:6–8.

Berger-Sweeney J, Heckers S, Mesulam MM, Wiley RG, Lappi D a, Sharma M (1994) Differential effects on spatial navigation of immunotoxin-induced cholinergic lesions of the medial septal area and nucleus basalis magnocellularis. J Neurosci 14:4507–4519.

Buzsaki G, Bickford RG, Ponomareff G, Thal LJ, Mandel R, Gage FH (1988) Nucleus basalis and thalamic control of neocortical activity in the freely moving rat. J Neurosci 8:4007–4026.

Chubykin AA, Roach EB, Bear MF, Shuler MGH (2013) A cholinergic mechanism for reward timing within primary visual cortex. Neuron 77:723–735.

Cohen JY, Haesler S, Vong L, Lowell BB, Uchida N (2012) Neuron-type-specific signals for reward and punishment in the ventral tegmental area. Nature 482:85–88.

Conner JM, Culberson A, Packowski C, Chiba AA, Tuszynski MH (2003) Lesions of the Basal forebrain cholinergic system impair task acquisition and abolish cortical plasticity associated with motor skill learning. Neuron 38:819–829.

Crouse RB, Kim K, Batchelor HM, Girardi EM, Kamaletdinova R, Chan J, Rajebhosale P, Pittenger ST, Role LW, Talmage DA, Jing M, Li Y, Gao X-B, Mineur YS, Picciotto MR (2020) Acetylcholine is released in the basolateral amygdala in response to predictors of reward and enhances the learning of cue-reward contingency. Elife 9:1–31.

Dalley JW, Theobald DE, Bouger P, Chudasama Y, Cardinal RN, Robbins TW (2004) Cortical cholinergic function and deficits in visual attentional performance in rats following 192 IgG-saporin-induced lesions of the medial prefrontal cortex. Cereb Cortex 14:922–932.

Damasio AR, Graff-Radford NR, Eslinger PJ, Damasio H, Kassell N (1985) Amnesia following basal forebrain lesions. Arch Neurol 42:263–271.

Disney AA, Aoki C, Hawken MJ (2007) Gain modulation by nicotine in macaque v1. Neuron 56:701–713.

Duque A, Balatoni B, Detari L, Zaborszky L (2000) EEG correlation of the discharge properties of identified neurons in the basal forebrain. J Neurophysiol 84:1627–1635.

Eggermann E, Kremer Y, Crochet S, Petersen CCH (2014) Cholinergic Signals in Mouse Barrel Cortex during Active Whisker Sensing. Cell Rep 9:1654–1661.

Endres DM, Schindelin JE (2003) A New Metric for Probability Distributions. IEEE Trans Inf Theory 49:1858–1860.

Eshel N, Tian J, Bukwich M, Uchida N (2016) Dopamine neurons share common response function for reward prediction error. Nat Neurosci 19:479–486.

Everitt BJ, Robbins TW (1997) Central cholinergic systems and cognition. Annu Rev Psychol 48:649–684.

Fine A, Hoyle C, Maclean CJ, Levatte TL, Baker HF, Ridley RM (1997) Learning impairments following injection of a selective cholinergic immunotoxin, ME20.4 IgG-saporin, into the basal nucleus of Meynert in monkeys. Neuroscience 81:331–343.

Franklin KB, Paxinos G (2007) The Mouse Brain in Stereotaxic Coordinates, Third Edition. Academic Press.

Fraser GW, Schwartz AB (2012) Recording from the same neurons chronically in motor cortex. J Neurophysiol 107:1970–1978.

Froemke RC, Merzenich MM, Schreiner CE (2007) A synaptic memory trace for cortical receptive field plasticity. Nature 450:425–429.

Gasselin C, Hohl B, Vernet A, Crochet S, Petersen CCH (2021) Cell-type-specific nicotinic input disinhibits mouse barrel cortex during active sensing. Neuron 109:778–787.e3.

Gershman SJ, Uchida N (2019) Believing in dopamine. Nat Rev Neurosci 20:703–714.

Gielow MR, Zaborszky L (2017) The Input-Output Relationship of the Cholinergic Basal Forebrain. Cell Rep 18:1817–1830.

Gombkoto P, Gielow M, Varsanyi P, Chavez C, Zaborszky L (2021) Contribution of the basal forebrain to corticocortical network interactions. Brain Struct Funct 226:1803–1821.

Gu Z, Yakel JL (2011) Timing-dependent septal cholinergic induction of dynamic hippocampal synaptic plasticity. Neuron 71:155–165.

Guo W, Robert B, Polley DB (2019) The Cholinergic Basal Forebrain Links Auditory Stimuli with Delayed Reinforcement to Support Learning. Neuron 103:1164–1177.e6.

Hangya B, Ranade SP, Lorenc M, Kepecs A (2015) Central Cholinergic Neurons Are Rapidly Recruited by Reinforcement Feedback. Cell 162:1155–1168.

Harris KD, Thiele A (2011) Cortical state and attention. Nat Rev Neurosci 12:509–523.

Harrison TC, Pinto L, Brock JR, Dan Y (2016) Calcium Imaging of Basal Forebrain Activity during Innate and Learned Behaviors. Front Neural Circuits 10:1–12.

Hasselmo ME, Sarter M (2011) Modes and models of forebrain cholinergic neuromodulation of cognition. Neuropsychopharmacology 36:52–73.

Hegedüs P, Heckenast J, Hangya B (2021a) Differential recruitment of ventral pallidal e-types by behaviorally salient stimuli during Pavlovian conditioning. iScience 24:102377.

Hegedüs P, Velencei A, Belval C-H de, Heckenast J, Hangya B (2021b) Training protocol for probabilistic Pavlovian conditioning in mice using an open-source head-fixed setup. STAR Protoc 2:100795.

Higley MJ, Gittis AH, Oldenburg I a, Balthasar N, Seal RP, Edwards RH, Lowell BB, Kreitzer AC, Sabatini BL (2011) Cholinergic interneurons mediate fast VGluT3-dependent glutamatergic transmission in the striatum. PLoS One 6:e19155.

Jiang L, Kundu S, Lederman JD, López-Hernández GY, Ballinger EC, Wang S, Talmage DA, Role LW (2016) Cholinergic Signaling Controls Conditioned Fear Behaviors and Enhances Plasticity of Cortical-Amygdala Circuits. Neuron 90:1057–1070.

Kilgard MP, Merzenich MM (1998) Cortical Map Reorganization Enabled by Nucleus Basalis Activity. Science (80-) 279:1714–1718.

Kim CK, Yang SJ, Pichamoorthy N, Young NP, Kauvar I, Jennings JH, Lerner TN, Berndt A, Lee SY, Ramakrishnan C, Davidson TJ, Inoue M, Bito H, Deisseroth K (2016) Simultaneous fast measurement of circuit dynamics at multiple sites across the mammalian brain. Nat Methods 13:325–328.

Kim HR, Malik AN, Mikhael JG, Bech P, Tsutsui-Kimura I, Sun F, Zhang Y, Li Y, Watabe-Uchida M, Gershman SJ, Uchida N (2020) A Unified Framework for Dopamine Signals across Timescales. Cell 183:1600–1616.e25.

Kvitsiani D, Ranade S, Hangya B, Taniguchi H, Huang JZ, Kepecs A (2013) Distinct behavioural and network correlates of two interneuron types in prefrontal cortex. Nature 498:363–366.

Lak A, Stauffer WR, Schultz W (2016) Dopamine neurons learn relative chosen value from probabilistic rewards. Elife 5:1–19.

Laszlovszky T, Schlingloff D, Hegedüs P, Freund TF, Gulyás A, Kepecs A, Hangya B (2020) Distinct synchronization, cortical coupling and behavioral function of two basal forebrain cholinergic neuron types. Nat Neurosci 23:992–1003.

Leão RN, Mikulovic S, Leão KE, Munguba H, Gezelius H, Enjin A, Patra K, Eriksson A, Loew LM, Tort ABL, Kullander K (2012) OLM interneurons differentially modulate CA3 and entorhinal inputs to hippocampal CA1 neurons. Nat Neurosci 15:1524–1530.

Lee MG, Hassani OK, Alonso A, Jones BE (2005) Cholinergic basal forebrain neurons burst with theta during waking and paradoxical sleep. J Neurosci 25:4365–4369.

Lerner TN, Shilyansky C, Davidson TJ, Evans KE, Beier KT, Zalocusky KA, Crow AK, Malenka RC, Luo L, Tomer R, Deisseroth K (2015) Intact-Brain Analyses Reveal Distinct Information Carried by SNc Dopamine Subcircuits. Cell 162:635–647.

Letzkus JJ, Wolff SBE, Meyer EMM, Tovote P, Courtin J, Herry C, Lüthi A (2011) A disinhibitory microcircuit for associative fear learning in the auditory cortex. Nature 480:331–335.

Lin S-C, Nicolelis M a L (2008) Neuronal ensemble bursting in the basal forebrain encodes salience irrespective of valence. Neuron 59:138–149.

Lin S, Brown RE, Hussain Shuler MG, Petersen CCH, Kepecs A (2015) Optogenetic Dissection of the Basal Forebrain Neuromodulatory Control of Cortical Activation, Plasticity, and Cognition. J Neurosci 35:13896–13903.

Liu C-H, Coleman JE, Davoudi H, Zhang K, Hussain Shuler MG (2015) Selective Activation of a Putative Reinforcement Signal Conditions Cued Interval Timing in Primary Visual Cortex. Curr Biol 25:1551– 1561.

Lovett-Barron M, Kaifosh P, Kheirbek M a, Danielson N, Zaremba JD, Reardon TR, Turi GF, Hen R, Zemelman B V, Losonczy A (2014) Dendritic inhibition in the hippocampus supports fear learning. Science (80-) 343:857–863.

Matsumoto M, Hikosaka O (2009) Two types of dopamine neuron distinctly convey positive and negative motivational signals. Nature 459:837–841.

McGaughy J, Dalley JW, Morrison CH, Everitt BJ, Robbins TW (2002) Selective behavioral and neurochemical effects of cholinergic lesions produced by intrabasalis infusions of 192 IgG-saporin on attentional performance in a five-choice serial reaction time task. J Neurosci 22:1905–1913.

McGaughy J, Koene R a, Eichenbaum H, Hasselmo ME (2005) Cholinergic deafferentation of the entorhinal cortex in rats impairs encoding of novel but not familiar stimuli in a delayed nonmatch-to-sample task. J Neurosci 25:10273–10281.

Menegas W, Akiti K, Amo R, Uchida N, Watabe-Uchida M (2018) Dopamine neurons projecting to the posterior striatum reinforce avoidance of threatening stimuli. Nat Neurosci 21:1421–1430.

Metherate R, Cox CL, Ashe JH (1992) Cellular bases of neocortical activation: modulation of neural oscillations by the nucleus basalis and endogenous acetylcholine. J Neurosci 12:4701–4711.

Najafi F, Giovannucci A, Wang SSH, Medina JF (2014) Coding of stimulus strength via analog calcium signals in Purkinje cell dendrites of awake mice. Elife 3:e03663.

Nelson A, Mooney R (2016) The Basal Forebrain and Motor Cortex Provide Convergent yet Distinct Movement-Related Inputs to the Auditory Cortex. Neuron 90:635–648.

Palacios-Filardo J, Mellor JR (2019) Neuromodulation of hippocampal long-term synaptic plasticity. Curr Opin Neurobiol 54:37–43.

Parikh V, Kozak R, Martinez V, Sarter M (2007) Prefrontal acetylcholine release controls cue detection on multiple timescales. Neuron 56:141–154.

Pi H-J, Hangya B, Kvitsiani D, Sanders JI, Huang ZJ, Kepecs A (2013) Cortical interneurons that specialize in disinhibitory control. Nature 503:521–524.

Pinto L, Goard MJ, Estandian D, Xu M, Kwan AC, Lee S-H, Harrison TC, Feng G, Dan Y (2013) Fast modulation of visual perception by basal forebrain cholinergic neurons. Nat Neurosci 16:1857– 1863.

Richardson RT, DeLong MR (1991) Electrophysiological studies of the functions of the nucleus basalis in primates. Adv Exp Med Biol 295:233–252.

Robert B, Kimchi EY, Watanabe Y, Chakoma T, Jing M, Li Y, Polley DB (2021) A functional topography within the cholinergic basal forebrain for encoding sensory cues and behavioral reinforcement outcomes. Elife 10:1–28.

Ross RS, McGaughy J, Eichenbaum H (2005) Acetylcholine in the orbitofrontal cortex is necessary for the acquisition of a socially transmitted food preference. Learn Mem 12:302–306.

Rouhani N, Niv Y (2021) Signed and unsigned reward prediction errors dynamically enhance learning and memory. Elife 10:1–28.

Rouhani N, Norman KA, Niv Y (2018) Dissociable effects of surprising rewards on learning and memory. J Exp Psychol Learn Mem Cogn 44:1430–1443.

Schmitzer-Torbert N, Jackson J, Henze D, Harris K, Redish a D (2005) Quantitative measures of cluster quality for use in extracellular recordings. Neuroscience 131:1–11.

Schultz W (2015) Neuronal Reward and Decision Signals: From Theories to Data. Physiol Rev 95:853– 951.

Schultz W, Dayan P, Montague PR (1997) A neural substrate of prediction and reward. Science 275:1593–1599.

Shuler MG, Bear MF (2006) Reward timing in the primary visual cortex. Science 311:1606–1609.

Siegle JH, López AC, Patel YA, Abramov K, Ohayon S, Voigts J (2017) Open Ephys: an open-source, plugin-based platform for multichannel electrophysiology. J Neural Eng 14:045003.

Solari N, Sviatkó K, Laszlovszky T, Hegedüs P, Hangya B (2018) Open Source Tools for Temporally Controlled Rodent Behavior Suitable for Electrophysiology and Optogenetic Manipulations. Front Syst Neurosci 12.

Stephenson-Jones M, Bravo-Rivera C, Ahrens S, Furlan A, Xiao X, Fernandes-Henriques C, Li B (2020) Opposing Contributions of GABAergic and Glutamatergic Ventral Pallidal Neurons to Motivational Behaviors. Neuron 105:921–933.e5.

Sviatkó K, Hangya B (2017) Monitoring the Right Collection: The Central Cholinergic Neurons as an Instructive Example. Front Neural Circuits 11:31.

Széll A, Martínez-Bellver S, Hegedüs P, Hangya B (2020) OPETH: Open Source Solution for Real-Time Peri-Event Time Histogram Based on Open Ephys. Front Neuroinform 14:1–19.

Teles-Grilo Ruivo LM, Baker KL, Conway MW, Kinsley PJ, Gilmour G, Phillips KG, Isaac JTR, Lowry JP, Mellor JR (2017) Coordinated Acetylcholine Release in Prefrontal Cortex and Hippocampus Is Associated with Arousal and Reward on Distinct Timescales. Cell Rep 18:905–917.

Tsutsui-Kimura I, Matsumoto H, Akiti K, Yamada MM, Uchida N, Watabe-Uchida M (2020) Distinct temporal difference error signals in dopamine axons in three regions of the striatum in a decision-making task. Elife 9:1–39.

Urban-Ciecko J, Jouhanneau JS, Myal SE, Poulet JFA, Barth AL (2018) Precisely Timed Nicotinic Activation Drives SST Inhibition in Neocortical Circuits. Neuron 97:611–625.e5.

Whitehouse PJ, Price DL, Struble RG, Clark AW, Coyle JT, Delon MR (1982) Alzheimer’s disease and senile dementia: loss of neurons in the basal forebrain. Science 215:1237–1239.

Wrenn CC, Wiley RG (1998) The behavioral functions of the cholinergic basal forebrain: lessons from 192 IgG-saporin. Int J Dev Neurosci 16:595–602.

Xu M, Chung S, Zhang S, Zhong P, Ma C, Chang W-C, Weissbourd B, Sakai N, Luo L, Nishino S, Dan Y (2015) Basal forebrain circuit for sleep-wake control. Nat Neurosci 18:1641–1647.

Zaborszky L, Csordas A, Mosca K, Kim J, Gielow MR, Vadasz C, Nadasdy Z (2013) Neurons in the Basal Forebrain Project to the Cortex in a Complex Topographic Organization that Reflects Corticocortical Connectivity Patterns: An Experimental Study Based on Retrograde Tracing and 3D Reconstruction. Cereb Cortex.

Zannone S, Brzosko Z, Paulsen O, Clopath C (2018) Acetylcholine-modulated plasticity in reward-driven navigation: a computational study. Sci Rep 8:9486.

Zhang H, Lin S-C, Nicolelis MAL (2011) A distinctive subpopulation of medial septal slow-firing neurons promote hippocampal activation and theta oscillations. J Neurophysiol 106:2749–2763.

Zhang K, Chen CD, Monosov IE (2019) Novelty, Salience, and Surprise Timing Are Signaled by Neurons in the Basal Forebrain. Curr Biol 29:134–142.e3.

